# Eco-tank Housing Maintains Wild-Type Microbiota and Rewilds the Laboratory Mouse Gut Microbiome to Restore Natural Immune Tone

**DOI:** 10.64898/2026.03.16.712136

**Authors:** Ti Lu, Zackary K. Dietz, Aaron C. Ericsson, William D. Picking, Wendy L. Picking

## Abstract

Laboratory mice housed under individually ventilated cage (IVC) conditions harbor simplified gut microbiota and immune phenotypes that diverge substantially from those shaped by environmental exposure, limiting translational relevance. To reintroduce controlled ecological complexity while maintaining biosafety and reproducibility, we developed the Eco-tank, a pathogen-monitored semi-natural housing system incorporating environmental substrates and dietary diversity. Longitudinal 16S rRNA sequencing revealed that even wild-caught *Mus musculus* rapidly lose microbial richness and predicted metabolic breadth under IVC housing. Eco-tank conditions stabilized diversity and preserved elements of wild-associated community structure during extended captivity. In parallel, standardized C57BL/6 mice housed in Eco-tanks underwent rewilding-like restructuring, with increased richness and community shifts toward a wild-associated configuration. Functional inference analyses indicated expansion of predicted pathways linked to short-chain fatty acid production, amino acid metabolism, and environmental substrate utilization. Eco-tank housing enhanced baseline resistance to pulmonary *Pseudomonas aeruginosa* (Pa) infection without compromising vaccine-induced protection, indicating that restoration of environmental microbial signals does not impair adaptive immunity. Together, these findings identify housing ecology as a dominant determinant of microbiome structure and functional potential. The Eco-tank provides a scalable and tractable framework for integrating environmental microbial complexity into laboratory models to better align preclinical immunology with ecologically conditioned immune systems.

**Importance:** Laboratory mice are foundational models for immunology, yet their specific pathogen-free rearing and housing environments impose ecological constraints that reshape the gut microbiome and immune tone. This study introduces a scalable, pathogen-monitored Eco-tank system that restores environmental microbial complexity while preserving experimental control. By demonstrating that housing ecology reshapes microbiome functional potential and modulates baseline immune resistance without compromising vaccine responsiveness, this work highlights environmental context as a critical experimental variable in preclinical immunology and offers a tractable framework for improving translational relevance.

## Introduction

The mammalian immune system develops and functions in constant interaction with the commensal microbiome, which provides microbial-derived signals that shape immune maturation, mucosal barrier function, and resistance to infection. Continuous exposure to microbial antigens and metabolites calibrates both innate and adaptive immune responses, particularly at barrier surfaces such as the gut and lung ^1–4^. Perturbations of host-microbiota interactions can alter susceptibility to pathogens and modify the magnitude and quality of vaccine-induced protection ^5–7^.

Despite the central role of the microbiome in immune regulation, most laboratory mouse studies are conducted under conventional specific pathogen-free (SPF) conditions that severely restrict environmental microbial exposure ^8^. In practice, these systems typically utilize individually ventilated microisolator cages (IVC) supplied with sterilized bedding, irradiated feed, and treated water, thereby minimizing environmental microbial exposure ^9^ (abbreviations used here are provided in Suppl. Table S1). Compared with wild mice and humans, IVC-housed mice harbor simplified and homogeneous microbiota and exhibit attenuated immune maturation ^10,11^. Many taxa and metabolic pathways prevalent in environmentally exposed and human-associated microbiomes are underrepresented in IVC-housed laboratory mice ^12,13^. Moreover, IVC mice have underdeveloped memory and effector T cell populations, fewer activated myeloid and NK cells, and reduced germinal center activity compared to wild mice ^11,14^. Researchers also found that lymphoid organs (e.g., Peyer’s patches, mesenteric lymph nodes) were smaller and less developed in IVC mice due to insufficient microbial-driven maturation signals ^15^. And serum immunoglobulin levels were lower in IVC mice, reflecting limited antigen exposure ^14^. These differences can exaggerate experimental phenotypes, including infection susceptibility and vaccine efficacy, raising concerns about the translational relevance of conventional laboratory models ^16–18^.

To address these limitations, multiple strategies have been developed to increase microbial exposure in laboratory mice, including co-housing with wild or pet store mice, transfer of wild microbiota, and outdoor rewilding enclosures ^16,19–23^. While these approaches have revealed important effects of environmental microbial exposure on immune complexity and disease resistance, they often introduce substantial variability, are difficult to standardize, and may lack rigorous pathogen monitoring. Seasonal effects, geographic heterogeneity, and uncontrolled pathogen introduction further limit reproducibility and biosafety, constraining broader adoption of these models ^24–26^. Consequently, the field lacks a housing system that captures key features of environmental microbial exposure while remaining standardized, pathogen-monitored, and readily transferable across research settings.

Here, we established and validated a pathogen-monitored, semi-natural housing system (Eco-tank) that integrated controlled environmental microbial input while maintaining experimental manageability. By combining soil, ecological, and dietary complexity within an indoor enclosure, the Eco-tank model preserved wild-associated microbiota structure and functional potential while operating under pathogen-monitored conditions that mitigate the biosafety and reproducibility limitations of outdoor rewilding approaches. Using wild-caught *Mus musculus* (*Mus*) and laboratory C57BL/6 mice, we showed that standard laboratory IVC housing rapidly erodes microbial diversity, whereas Eco-tank housing maintained ecological proximity to wild microbiota and revealed infection phenotypes masked under standard laboratory conditions.

## Methods

### Eco-tank (Semi-Natural Stock Tank) Setup

A subset of experiments was conducted in semi-naturalistic enclosures (Eco-tanks) constructed from galvanized steel stock tanks (2 × 2 × 6 ft; CountyLine, Tractor Supply Co., Columbia, MO) and housed indoors under ambient environmental conditions. Tanks were covered with custom-fitted, mesh-screened lids secured in painted wooden frames to prevent animal escape while allowing airflow. Animals were maintained under a natural light/dark cycle with ambient temperature (20-22°C) and humidity (40-60%). To simulate natural substrate and facilitate drainage, each tank was layered with approximately 1 inch of river rock (1.5 cu ft/tank; Quikrete, ACE Hardware, Columbia, MO), overlaid with 2 inches of native soil collected from Boone County, Missouri (10-15 gallons/tank). Water was provided *ad libitum* via hanging water bottles.

Mice were fed a custom diet (**Naturalistic Diet**) consisting of 95% wild birdseed (Wagner’s Eastern Regional Blend Deluxe Wild Bird Food, USA) and 5% dried mealworms (Coohgrubs Dried Mealworms Poultry & Bird Treats, USA), delivered in gravity feeders placed directly on the soil surface. Standard laboratory chow was not provided. Animals were allowed to breed freely within each enclosure to permit natural population dynamics. Environmental enrichment included nesting material (Bed-r’Nest) and naturalistic features such as sticks and PVC piping (ACE Hardware) to support burrowing, provide cover, and allow exploratory behaviors (**Figure 1**). All out-of-laboratory materials were obtained in batches from a single consistent source and subjected to pathogen screening prior to use to ensure safety and reproducibility across enclosures.

**Figure 1.**
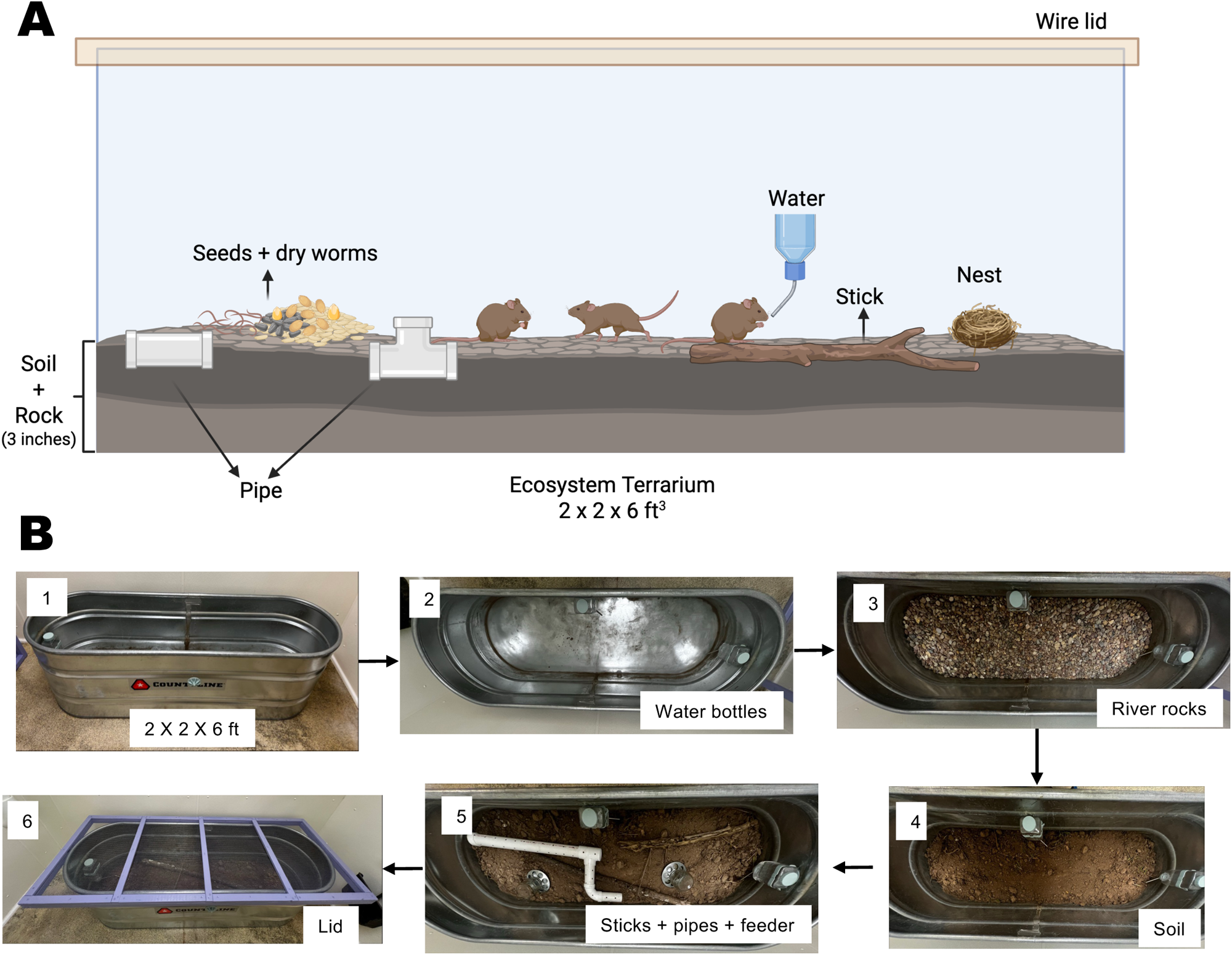
Semi-naturalistic Eco-tank enclosure for wild-derived mouse studies. (**A**) Schematic of the Eco-tank housing system illustrating enclosure dimensions (2 × 2 × 6 ft), layered substrate (river rock and soil), food and water placement, nesting material, and environmental enrichment. Tanks were covered with a mesh-screened lid to prevent escape while allowing ventilation. Figure was generated by using BioRender. (**B**) Stepwise assembly of the Eco-tank system, including placement of water bottles, river rock drainage layer, soil substrate, enrichment materials (sticks and PVC pipes), and the final secured lid.

### Animals and Housing

All animal protocols were reviewed and approved by the University of Missouri Institutional Animal Care and Use Committee Practices (protocols 38241 and 65063). Wild *Mus musculus* (n = 24) were live-trapped on the University of Missouri-Columbia campus and designated as wild-state condition (**Wild**) at the time of capture. Immediately following capture, animals were transported to the laboratory and examined by veterinary staff to confirm species identity based on morphological criteria and to assess overall health status. Fecal pellets and pelt swabs were submitted to a diagnostic laboratory (IDEXX BioAnalytics, Columbia, MO) for pathogen screening using the Opti-XXpress/EDx Mouse Global PCR panel, which tests for a defined set of bacterial, viral, and parasitic agents including several zoonotic pathogens. No human pathogens included in the diagnostic panel were detected. Detected microorganisms consisted primarily of commensal or rodent-associated taxa commonly reported in wild mouse populations. Detailed screening results are provided in Suppl. Table S2. Only animals verified as *Mus musculus* and deemed healthy were included in subsequent breeding and experimental procedures.

During veterinary screening, mice were housed under standard barrier laboratory conditions in individually ventilated microisolator cages (**IVC**) on ventilated racks ^9^. Cages were supplied with autoclaved bedding, irradiated standard rodent chow, and autoclaved drinking water provided ad libitum ^27^. Cage changes were performed under biosafety cabinet conditions using sterile technique. This 14-day acclimation period generated a laboratory-adapted “IVC-like” state (**IVC-like cage**; Suppl. Figure S1A). Then wild-caught *Mus* were randomly assigned to one of four treatment groups (n = 6 per group; 4 females and 2 males per group) to assess whether environmental and dietary enrichment could mitigate microbiota loss. (1) Chow-only (**IVC-like cage**): Mice continued standard chow diet with regular laboratory bedding and no additional enrichment. (2) Chow + Soil (**Soil bedding**): Mice were housed with native soil collected from Boone County, MO added to regular bedding at a 1:2 soil-to-bedding ratio, while continuing standard chow. (3) Chow + **Naturalistic Diet**: Mice received a mixed diet consisting of standard chow supplemented with a Naturalistic Diet (95:5 birdseed-to-mealworm ratio) while housed on regular bedding. (4) Chow + Soil + Naturalistic Diet (**Semi-natural cage**): Mice received both native soil bedding and standard chow supplemented with Naturalistic Diet. These conditions were maintained for 10 days (Day 14-24). On Day 24, mice from Groups 2-4 were ear-tagged and transferred into a large shared Eco-tank system (Eco-tank 1) designed to provide sustained environmental complexity and microbial exposure (Suppl. Figure S1A).

All cages and the Eco-tank system were maintained in the same animal room under identical ambient environmental parameters (as described above). Prior to sample collection, mice within each experimental group were housed together in a single large enclosure (one cage per group), allowing shared environmental exposure and microbial exchange within each group. After sample collection, animals were reorganized into additional cages as necessary to reduce crowding. Because animals experienced the same enclosure environment prior to sampling, microbiome analyses focused on group-level differences rather than cage-level variation. To assess the reproducibility of Eco-tank mediated preservation of wild gut microbiota, a second Eco-tank (Eco-tank 2) was established in the same animal room with materials and batches from the same resources and maintained under identical environmental parameters as Eco-tank 1. Wild-caught *Mus* that had remained in IVC-like cages on standard chow diet through Day 52 (Group 1) were transferred into Eco-tank 2. Both traditional IVC housing and eco-tank environments were maintained under institutional sentinel monitoring for excluded human pathogens as described above.

To test whether Eco-tank housing can shape the gut microbiome in standard laboratory mice, commercially sourced C57BL/6 mice (n = 10; C57BL/6NCrl; Charles River Laboratories, Wilmington, MA) were used. Founder animals were bred and maintained under IVC conditions. At weaning, F1 offspring were assigned to one of three housing conditions: (1) IVC-like cage; (2) Semi-natural cage; (3) Eco-tank. Each group consisted of comparable numbers of males and females (Eco-tank: 10 females, 2 males; IVC and semi-natural cages: 5 females, 5 males per group). Housing conditions were initiated at weaning and maintained 28 days prior to microbiota sampling. Eco-tank and cage housing for C57BL/6 mice was conducted under environmental conditions matched to those used for wild *Mus*. All housing systems were located within the same animal facility and maintained under identical ambient environmental conditions (as described above). Experiments involving C57BL/6 mice were performed in a separate sub-room within the same laboratory area to prevent cross-contamination while preserving equivalent environmental parameters.

### Sampling

During the experiment, wild-caught *Mus* and C57BL/6 bred naturally within the housing systems, including in the Eco-tank environment. As a result, fecal samples collected at later time points, including Day 52, may include contributions from offspring born during the experimental period. Offspring (> 21 days post birth) were not separated from parental animals and were exposed continuously to the same housing, dietary, and environmental conditions as the founding adults. Fecal samples from wild *Mus* were collected at four time points: Day 0 (from live traps within 12 h of capture), Day 14 (after maintenance in standard laboratory cages on a standard chow diet), Day 24 (following 10 days in treatment groups 2-4), and Day 52 (from treatment 1 and Eco-tank enclosures, 28 days post-transfer). While all fecal samples from C57BL/6 were collected at the same timepoint (28 days post environmental transfer). Environmental soil used in Eco-tank housing was sampled from four independent containers for 16S rRNA gene sequencing to characterize the microbial composition of the environmental input.

Samples were stored at -80°C until DNA isolation was performed. As animals were not individually tracked, fecal pellets were collected at the group level from the collection container or traps within each treatment or enclosure. Frozen samples were shipped directly on dry ice to the University of Missouri Metagenomics Core (MUMC, Columbia, MO) for DNA extraction, preparation, and plating within one week.

### DNA extraction

DNA was extracted as previously described ^28^. DNA was isolated using QIAamp PowerFecal Pro DNA kits (Qiagen) according to the manufacturer’s instructions, with a modification to the sample disruption step. Instead of the vortex adapter recommended in the protocol, bead-beating was performed using a TissueLyser II (Qiagen, Venlo, Netherlands) at 30 Hz for 10 minutes. Subsequent steps followed the standard protocol, and DNA was eluted in 100 µL of Qiagen elution buffer. DNA yield was quantified by fluorometric analysis using a Qubit 2.0 instrument (Invitrogen, Carlsbad, CA) with the Quant-iT BR double-stranded DNA assay kit, after which samples were normalized to a consistent concentration and volume ^29^.

### 16S rRNA library preparation and sequencing

Library preparation and sequencing were performed at the MU Genomics Technology Core as previously described ^28^. The bacterial community was profiled by amplifying the V4 hypervariable region of the 16S rRNA gene using the universal primer pair U515F and 806R, each incorporating Illumina adapter sequences ^30,31^. Dual-index barcodes were included on both forward and reverse primers. PCR amplification was carried out in 50 µL reaction volumes containing 100 ng of extracted fecal DNA, 0.2 µM of each primer, 200 µM of each dNTP, and 1 U of Phusion High-Fidelity DNA Polymerase (Thermo Fisher Scientific). Thermal cycling consisted of an initial denaturation at 98°C for 3 minutes, followed by 25 cycles of denaturation at 98°C for 15 seconds, annealing at 50°C for 30 seconds, and extension at 72 °C for 30 seconds, with a final extension at 72°C for 7 minutes. Following amplification, 5 µL from each reaction was combined into a single pool. The pooled amplicons were purified using Axygen Axyprep MagPCR magnetic beads at a 1:1 bead-to-sample volume ratio and incubated at room temperature for 15 minutes. Beads were subsequently washed multiple times with 80% ethanol, air-dried, and DNA was eluted in 32.5 µL of EB buffer (Qiagen). After a brief incubation at room temperature, samples were placed on a magnetic rack for five minutes to separate beads from eluate. The final amplicon library was assessed using an Advanced Analytical Fragment Analyzer, quantified with the Quant-iT high-sensitivity dsDNA assay, and diluted according to Illumina guidelines prior to sequencing on a MiSeq i100 platform using 2 × 300 bp paired-end chemistry.

### Informatics analysis

DNA sequences were assembled and annotated at the MU Bioinformatics and Analytics Core as previously described ^28^. Primer sequences were designed to correspond to the 5′ ends of both forward and reverse reads. Primer trimming was performed using Cutadapt (v2.6) ^32^, first removing the forward primer from the 5′ end of the forward read. If detected, the reverse complement of the reverse primer was also identified within the forward read and removed together with any downstream bases, allowing trimming at both ends in cases where the insert length was shorter than the expected amplicon. An analogous strategy was applied to the reverse reads, with primer roles reversed. Read pairs were discarded if either read failed to match a 5′ primer, permitting a maximum error rate of 0.1. Primer removal additionally required a minimum three-base pair overlap with the 3′ end of the primer sequence. Denoising, sequence de-replication, and amplicon sequence variant (ASV) inference were performed using the DADA2 ^33^ plugin (v1.10.0) within QIIME2 ^34^. Processing parameters included truncation of both forward and reverse reads to 150 bp, exclusion of reads with expected error values exceeding 2.0, and identification and removal of chimeric sequences using the consensus approach. Taxonomic classification of the resulting ASVs was conducted against the SILVA v132 ^35^ database (99% identity, non-redundant SSU) using the classify-sklearn algorithm.

### Microbiome Community Analysis

The original ASV count table (unfiltered and unrarefied) was used for diversity calculations and community-level analyses ^36^. Alpha diversity analyses were performed in R (v4.5.0) ^37,38^ using the vegan package (v2.7-2) ^39^. For Shannon diversity, ASV counts were converted to relative abundances using total sum scaling (TSS) ^40^. Shannon diversity was calculated using the diversity function ^27^. Chao1 richness estimates were computed from raw count data using the estimateR function, and the S.chao1 metric was extracted for downstream analysis ^27^. Beta diversity analyses were performed in R with the vegdist function in the vegan package in R ^39^, with relative abundance calculated by TSS. Bray-Curtis dissimilarity was computed on relative abundance data, and Jaccard distance was calculated on presence/absence-transformed ASV tables (ASV > 0). Ordination was visualized using principal coordinates analysis (PCoA) ^27,39^. The percentage of variance explained by each axis was calculated from eigenvalues and displayed on ordination plots. Ellipses represent 95% confidence intervals. Environmental soil samples were excluded from statistical testing but included in separate ordination visualizations for ecological context. Distance-to-wild was defined as the mean Bray-Curtis dissimilarity between each sample and all wild reference samples. Housing conditions were encoded as an ordered ecological exposure gradient (IVC-like < Semi-natural < Eco-tank < Wild) for directional analyses.

### Taxonomic abundance profiling

ASV tables and taxonomic assignments generated from 16S rRNA gene sequencing were aggregated at the phylum level prior to downstream analysis ^34,41^. Relative abundance was calculated by normalizing counts to total sequencing depth per sample^27^. Phylum-level abundance profiles were analyzed in R using the dplyr ^42^, tidyr ^43^, and ggplot2 ^44^ packages. To visualize taxonomic composition, the top 16 most abundant phyla across all samples were identified based on mean relative abundance and retained for downstream analyses, while remaining taxa were grouped as “Other.” Group means and individual sample distributions were visualized using stacked bar plots generated in ggplot2. Directional responses of dominant phyla along the ecological exposure gradient were visualized by plotting Spearman correlation coefficients (ρ) between phylum-level relative abundance and ordered housing conditions in R.

### Functional prediction analysis

Functional potential of the gut microbiota was predicted using TAX4FUN2 ^45^, implemented through MicrobiomeAnalyst 2.0 ^46,47^. Predictions were generated from ASV-level taxonomic assignments using both rarefied and TSS scaled ASV tables as input. Predicted functional profiles were summarized as KEGG Ortholog (KO) abundances ^48^. For downstream analyses, KO abundance tables were exported and processed in R ^37^. Functional abundances were normalized to relative abundance per sample and log10-transformed (log10[x + 1e-6]) prior to multivariate analysis ^45^. Features with zero variance across samples were removed. Principal component analysis (PCA) was performed using prcomp on log-transformed, scaled functional abundance matrices ^49^. The percentage of variance explained by each principal component was calculated from eigenvalues.

Functional pathways associated with housing condition were first identified using the functional association analysis module in MicrobiomeAnalyst 2.0 ^46,47^, which applies a global test framework to detect KEGG pathways whose collective gene abundances are significantly associated with the experimental factor (housing condition). All uploaded taxa and KEGG Orthologs were included in the metabolic network origin. For pathways identified as significantly associated with housing condition, pathway-level abundances were extracted and used to calculate log2 fold changes among each comparison ^47^. For functional analyses, Eco-tank replicate cohorts (Eco-tank 1 and Eco-tank 2) were combined into a single Eco-tank group, and early (14 days) and late (52 days) IVC samples were combined into a single IVC-like group. Although minor differences were observed between Eco-tank replicates at the ASV-level beta diversity, predicted functional profiles did not exhibit systematic separation between replicates. Therefore, samples were pooled to increase statistical power and to evaluate housing-dependent functional effects at the group level.

### Protein preparation, Vaccine formulation and C57BL/6NCrl mice immunization

L-PaF is a fusion protein of the *Pseudomonas aeruginosa* type III secretion system proteins PcrV and PopB possessing the LTA1 subunit of enterotoxigenic *Escherichia coli* heat-labile toxin located at its N-terminus ^50^. L-PaF was produced, purified, and quality-controlled as previously described ^50–52^. Briefly, the recombinant proteins were expressed in *E. coli* BL21(DE3), purified by affinity and size-exclusion chromatography, and formulated under endotoxin-controlled conditions. Endotoxin levels were confirmed to be <5 EU/mg protein using the Endosafe nexgen-PTS system (Charles River Laboratories, Wilmington, MA).

L-PaF was formulated with a squalene-based oil-in-water emulsion (ME) admixed with BECC438 (a nontoxic lipid-A analogue and TLR-4 agonist that provides additional adjuvant activity) as previously reported ^53,54^. The ME/L-PaF/BECC438 formulation is now called the **B**roadly-protective **R**ecombinant **A**ntigen **V**accine **E**nabled vaccine (**BRAVE.0**), however, because this study is focused on the host microbiome, the vaccine was only used to assess immune responsiveness to and protective efficacy against *P. aeruginosa*. C57BL/6 mice used for vaccination and infection experiments were independent individuals from the same housing cohorts and were not previously subjected to microbiome sampling. Animals were allowed to equilibrate (14 days) to their respective housing conditions (semi-natural cages: n = 8-9 per group; Eco-tank: n=12 per group) prior to immunization. Mice were then immunized intranasally under isoflurane anesthesia with recombinant L-PaF (1 μg/mouse) formulated in BECC/ME (hereafter referred to simply as BRAVE.0) ^35^. Immunizations were administered on days 0, 14, and 28, as indicated in the experimental timeline (Suppl. Figure S1C). Control animals received phosphate-buffered saline (PBS) alone following the same schedule. Mice remained in their assigned housing conditions for the duration of the immunization period.

### *Pseudomonas aeruginosa* (Pa) preparation and animal infection

For *in vivo* infection experiments, *P. aeruginosa* clinical strain mPA08-31 was plated from a frozen stock onto *Pseudomonas* isolation agar (PIA), followed by overnight incubation at 37°C ^35^. Single colonies were used to inoculate LB broth and cultured overnight at 37°C with shaking (200 rpm). The following day, bacterial cultures were diluted into fresh LB and grown to OD_600_ around 0.3. Cells were harvested by centrifugation (4000 rpm), washed once with sterile PBS, and resuspended in PBS. Bacterial suspensions were adjusted to the indicated concentrations for infection based on optical density and confirmed by plating. On day 28 post last vaccination, naïve and vaccinated C57BL/6 mice from semi-natural cages (n = 8-9 per group) or from Eco-tank (n = 12 per group) were anesthetized with isoflurane and intranasally challenged with 4 × 10^7^ CFU/30 μL; health and weight were monitored twice daily for one day, after which mice were euthanized, lungs harvested, and bacterial burden quantified via CFU enumeration on PIA plates ^35^.

### Statistics

All statistical analyses were performed in R (v4.5.0) ^37,38^ (unless otherwise specified) using the packages vegan ^39^, ggplot2 ^44^, dplyr ^42^, rstatix ^55^, and purrr ^56^. For comparisons between two groups, statistical significance was assessed using the Wilcoxon rank-sum test ^55^. For analyses involving more than two groups, differences were evaluated using the Kruskal-Wallis test, followed by pairwise Wilcoxon rank-sum tests with Benjamini-Hochberg false discovery rate (FDR) correction where applicable ^55^. All tests were two-sided. Microbial community differences were assessed using Bray-Curtis and Jaccard dissimilarity metrics calculated using the vegdist function in the vegan R package ^39^. Group differences in community composition were tested using permutational multivariate analysis of variance (PERMANOVA) implemented in adonis2 with 999 permutations, and R^2^ values were reported as effect sizes ^57^. Homogeneity of multivariate dispersion was evaluated using PERMDISP (betadisper) prior to interpretation ^58^. For multiple-group comparisons, pairwise PERMANOVA analyses were performed with FDR correction ^57^. Extracted R^2^ values were visualized as heatmaps, with color intensity representing effect size magnitude and statistical significance indicated by asterisks ^44,59^.

To quantify ecological convergence toward the wild microbiota, distance-to-wild was defined as the mean Bray-Curtis dissimilarity between each sample and all wild reference samples ^60^. Housing conditions were encoded as an ordered ecological exposure gradient (IVC-like < Semi-natural < Eco-tank < Wild). Differences among groups were tested using Kruskal-Wallis tests with pairwise Wilcoxon comparisons, and associations along the ecological gradient were evaluated using Spearman correlation and linear regression ^61^. Genotype-dependent responses to housing conditions were evaluated using a two-factor PERMANOVA model (Genotype × Housing) applied to mouse samples only (excluding wild-state animals), with marginal effects estimated using the by = “margin” argument in adonis2 ^57,61^.

Multiple testing correction was performed using the Benjamini-Hochberg false discovery rate (FDR) procedure in predicted function analysis, including functional association and differential pathway analyses ^46^. Differential pathway enrichment was visualized using volcano plots [x: log2 fold; y: -log(FDR)] and bar plots [x: log2 fold; y: Top 10 KEGG pathway with -log(FDR) > 2] generated in GraphPad Prism (Version 10.6.1). For *P. aeruginosa* lung challenge experiments, bacterial burdens (CFU) were compared using the Mann-Whitney test. Statistical significance was defined as *p* < 0.05 unless otherwise indicated ^50^.

## Results

### Rapid loss of wild gut microbiota diversity following laboratory acclimation

Preservation of native gut microbial communities remains a major challenge in studies using wild-caught animals. To determine how short-term laboratory housing alters microbiome structure, we compared ASV-level community composition in wild *Mus musculus* (*Mus*) at the time of capture (Wild) and after 14 days of housing under standard laboratory conditions (IVC housing). Wild mice at capture exhibited significantly higher alpha diversity compared to IVC-housed animals (**Figure 2A**). Both Chao1 richness (**Figure 2A** left) and Shannon diversity (**Figure 2A** right) were reduced following laboratory acclimation (Wilcoxon rank-sum test, *p* = 0.014 for both metrics), indicating rapid loss of microbial richness and evenness within two weeks of standardized housing.

**Figure 2.**
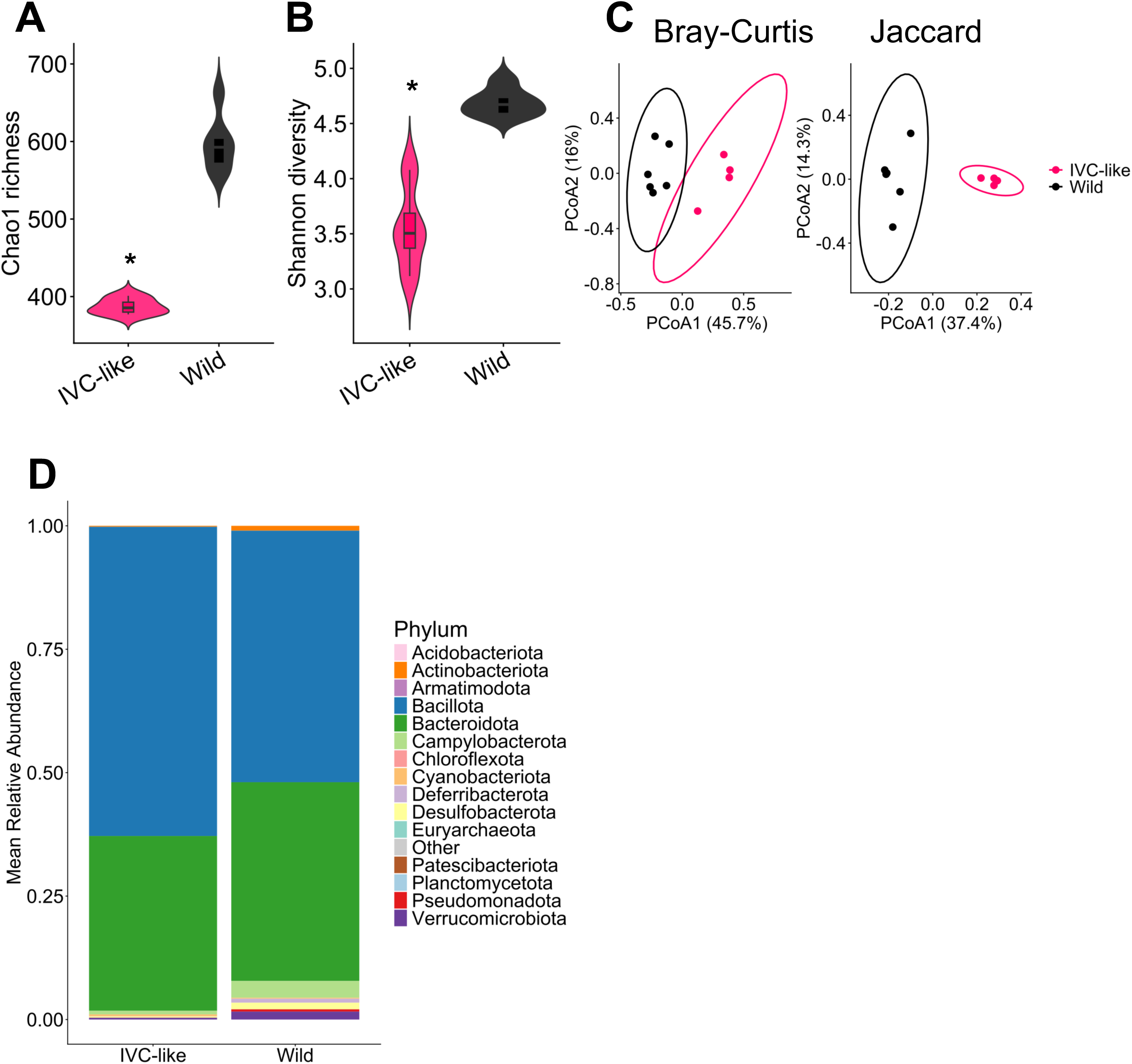
Rapid loss of wild gut microbiota diversity and restructuring following laboratory acclimation. Alpha diversity of gut microbiota at the ASV level in wild *Mus musculus* at capture (Wild: black) and after 14 days of standard laboratory housing (IVC-like: pink). Chao1 richness (**A**) and Shannon diversity (**B**) are shown as violin plots with embedded boxplots. Both richness and diversity were significantly reduced following laboratory acclimation (Wilcoxon rank-sum test, *p* = 0.014). (**C**) Principal coordinates analysis (PCoA) of ASV-level community structure based on Bray-Curtis (left) and Jaccard (right) dissimilarities. Each point represents an individual mouse; ellipses indicate 95% confidence intervals. Bray-Curtis analysis demonstrated significant compositional separation between Wild and IVC-like groups (PERMANOVA R^2^ = 0.421, F = 5.82, *p* = 0.003), with no significant difference in dispersion (PERMDISP *p* = 0.138). Jaccard analysis similarly showed significant separation (PERMANOVA R^2^ = 0.367, F = 4.63, p = 0.005), although dispersion differed modestly between groups (PERMDISP *p* = 0.024). (**D**) Phylum-level taxonomic composition of gut microbiota in wild and IVC-like mice. Bars represent mean relative abundance averaged across samples within each group, illustrating compositional shifts associated with laboratory housing. Phylum-level taxonomy was updated to conform with GTDB nomenclature ^84^.

ASV-level beta diversity analysis demonstrated marked restructuring of the microbial community. Bray-Curtis dissimilarity revealed clear separation between Wild and IVC-like groups (PERMANOVA R^2^ = 0.421, F = 5.82, *p* = 0.003; **Figure 2B** left). Importantly, homogeneity of dispersion did not differ significantly between groups for Bray-Curtis distances (PERMDISP *p* = 0.138), indicating that the observed separation reflects a shift in community centroid rather than differences in within-group variability. Analysis using Jaccard distance (presence/absence) similarly demonstrated significant compositional separation (PERMANOVA R^2^ = 0.367, F = 4.63, *p* = 0.005). Nevertheless, dispersion differed modestly between groups for Jaccard distances (PERMDISP *p* = 0.024), suggesting that both compositional shifts and altered community variability contributed to separation under presence-absence metrics.

At the phylum level, laboratory acclimation was accompanied by substantial compositional restructuring (**Figure 2C**). Wild-state microbiota displayed a taxonomically heterogeneous profile characterized by representation across multiple phyla, including *Patescibacteriota* (*Patescibacteria*), *Psdudomonadota*, *Verrucomicrobiota*, *Deferribacterota*, *Desulfobacterota*, and *Chloroflexota*. In contrast, IVC-like housing was associated with relative expansion of *Bacillota* and modest increases in *Cyanobacteriota*, resulting in a microbiome dominated by fewer high-abundance phyla. These data demonstrate that short-term exposure to standardized laboratory housing rapidly reduced alpha diversity and induced substantial ASV-level compositional remodeling. Notably, these extensive taxonomic shifts occurred within only two weeks of laboratory acclimation, indicating that wild-associated gut microbiota were extremely sensitive to standardized housing and diet and undergo rapid remodeling on a timescale that would be substantially shorter than many commonly used immunological interventions.

### Eco-tank housing stabilizes microbial diversity but does not fully recapitulate wild community structure

We next evaluated whether environmental and dietary enrichment could mitigate the progressive loss of wild-associated gut microbiota during extended captivity. Global analysis across housing conditions revealed significant differences in both ASV-level alpha diversity (Kruskal-Wallis: Chao1 *p* = 2.04 × 10^-7^, **Figure 3A**; Shannon *p* = 2.06 × 10^-6^, **Figure 3B**) and beta diversity (Bray-Curtis: PERMANOVA R^2^ = 0.438, F = 7.64, *p* = 0.001; **Figure 3C** left), with no significant difference in dispersion among groups (PERMDISP *p* = 0.129), indicating true compositional restructuring rather than variance artifacts. Similar patterns were observed using Jaccard dissimilarity (**Figure 3C** right; PERMANOVA R^2^ = 0.3792, *p* = 0.001; PERMDISP *p* = 0.114). Consistent with earlier observations, mice maintained under IVC-like chow-only conditions exhibited persistently reduced richness and diversity relative to wild-state animals. Soil bedding or wild dietary supplementation alone produced only modest improvements, and neither intervention restored diversity to wild-state levels (Suppl. Table S3). In contrast, Eco-tank housing substantially stabilized alpha diversity over prolonged captivity (Suppl. Table S3). By Day 52, Chao1 richness and Shannon diversity in Eco-tank mice were not significantly different from wild-state animals (pairwise Wilcoxon, FDR-adjusted *p* = 0.953 and *p* = 0.729, respectively), indicating near-complete restoration of overall diversity metrics.

**Figure 3.**
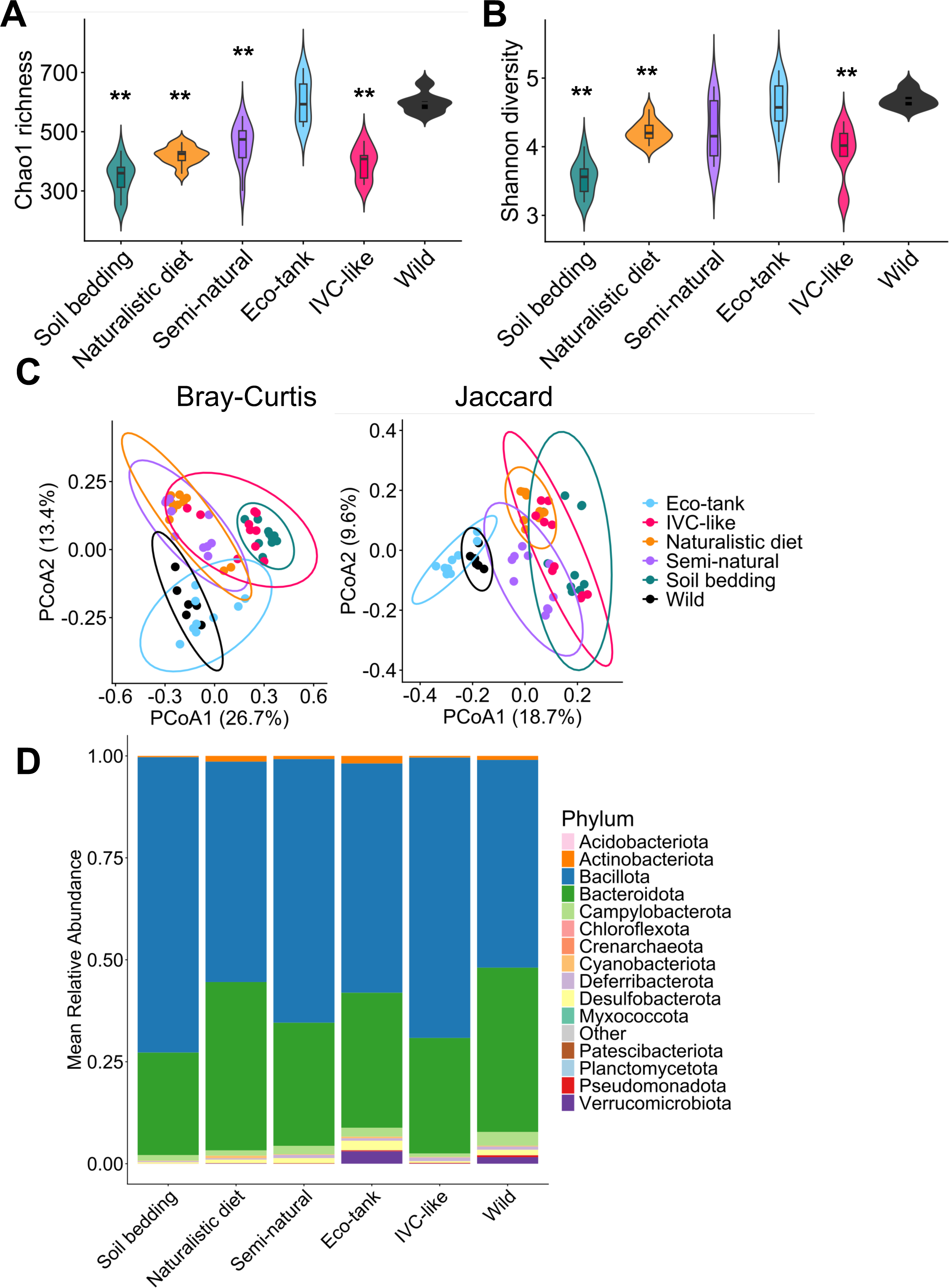
Eco-tank housing preserves wild-like gut microbiota diversity and community structure during prolonged captivity. Alpha diversity of gut microbiota across housing and dietary conditions, assessed by the Chao1 richness (**A**) and Shannon diversity (**B**) index. Groups include freshly captured wild mice (Wild; black), IVC-housed (pink), soil bedding (green), Naturalistic diet (yellow), semi-natural cages (purple), and Eco-tank housing (blue). Violin plots indicate distribution. Group *p* value by Kruskal-Wallis test with Dunn’s post hoc correction (Shannon: *p* < 0.001; Chao1: *p* < 0.001). ** *p* < 0.01 compared to Wild based on Pairwise comparisons using Wilcoxon rank sum test with continuity correction. (**C**) PCoA of gut microbial community composition based on Bray-Curtis dissimilarity (left; R^2^ = 0.43814, F = 7.642, *p* = 0.001) or Jaccard distance (right; R^2^ = 0.37924, F = 5.9872, *p* = 0.001). Each point represents individual fecal sample, colored by house conditions. Homogeneity of dispersion did not differ between tanks (PERMDISP Bray-Curtis *p* = 0.83; Jaccard *p* = 0.155), indicating that observed differences reflect compositional shifts rather than unequal within-group variability. Percent variance explained is shown on axes. Group differences were tested by PERMANOVA (999 permutations). Ellipses indicate 95% confidence intervals. (**D**) Phylum-level taxonomic composition of gut microbiota across experimental conditions at the indicated time points. Bars represent mean relative abundance across samples within each group. Phylum-level taxonomy was updated to conform with GTDB nomenclature ^84^. Statistical details of this figure are provided in Suppl. Table S3.

ASV-level community composition further supported that Eco-tank housing restores diversity but not identity. Pairwise PERMANOVA comparisons demonstrated that Eco-tank microbiota remained significantly different from wild-state communities (R^2^ = 0.237, FDR-adjusted *p* = 0.001; Suppl. Table S3), although the magnitude of separation was smaller than that observed for soil bedding, naturalistic diet alone, semi-natural cages, or IVC-like housing (R^2^ = 0.27-0.48; Suppl. Table S3 and Suppl. Figure S2A), supporting close similarity of Eco-tank to the wild state. Phylum-level profiling supported this intermediate positioning (**Figure 3D**; individual sample data in Suppl. Figure S2B). IVC-like housing was characterized by dominance of *Bacillota* and contraction of multiple low-abundance phyla. Soil bedding, dietary supplementation, and semi-natural cages showed limited ability to rebalance this architecture. Eco-tank mice, however, retained a broader phylum-level distribution more closely resembling wild-state animals, including representation of *Chloroflexota* (*p* = 0.47), *Verrucomicrobiota* (*p* = 0.65), and other low-abundance lineages, though subtle differences remained (Suppl. Figure S2C).

To test whether ecological exposure drives progressive convergence toward wild microbiota, we assigned housing conditions an ordered exposure score (IVC-like < Semi-natural < Eco-tank < Wild; **Figure 4A**). Spearman correlation analysis revealed a significant negative association between ecological exposure and Bray-Curtis distance to wild microbiota (ρ = −0.48, *p* = 0.003; **Figure 4B**), indicating monotonic convergence. Linear modeling confirmed this relationship (β = −0.031 per exposure level, *p* = 0.0087; R^2^ = 0.19). Phylum-level analysis revealed structured directional responses along the ecological exposure gradient (**Figure 4C**). *Verrucomicrobiota* (ρ = 0.71, FDR *p* = 2.6 × 10^-5^) and *Desulfobacterota* (ρ = 0.70, FDR *p* = 2.6 × 10^-5^) exhibited strong positive associations with ecological exposure, indicating enrichment toward wild-like states. In contrast, *Bacillota* abundance decreased significantly with increasing exposure (ρ = −0.63, FDR *p* = 3.1 × 10^-4^), consistent with reduced dominance of IVC-associated taxa. Together, these findings demonstrate exposure-dependent taxonomic restructuring that establishes a graded ecological restoration trajectory, indicating that sustained environmental complexity in the eco-tank promotes substantial convergence toward wild-associated community structure without fully recapitulating the wild microbiome.

**Figure 4.**
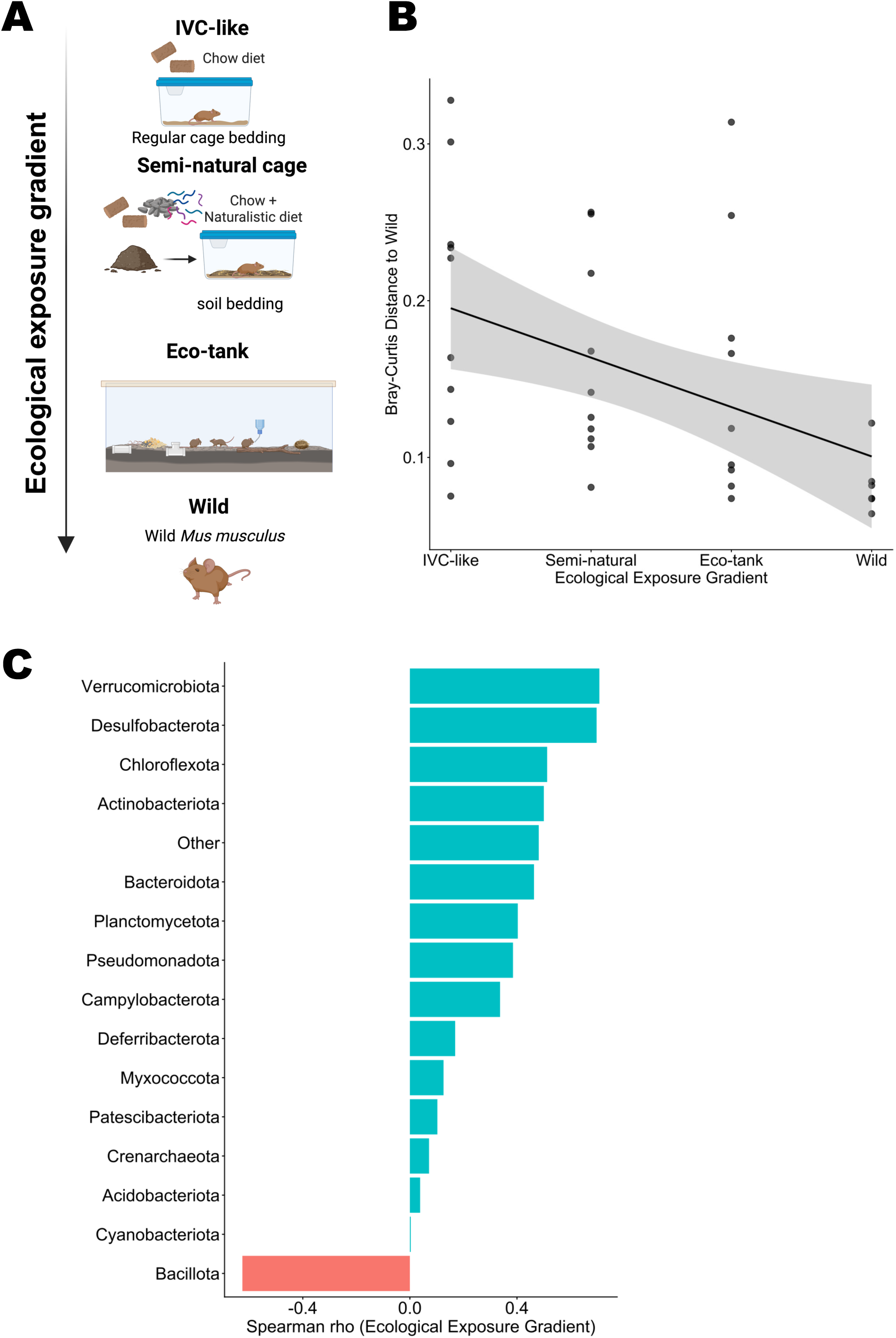
Ecological Exposure Drives Progressive Convergence Toward Wild Microbiota. (**A**) Schematic overview of the housing conditions used to increase environmental microbial exposure. This design establishes an ecological gradient from IVC housing to environmentally enriched conditions, enabling assessment of microbiome convergence toward the wild state. (**B**) Bray-Curtis distance of individual samples to the wild microbiota centroid plotted across an ordered ecological exposure gradient (IVC-like < Semi-natural < Eco-tank < Wild). Points represent individual fecal samples; line indicates linear regression with 95% confidence interval shading. Increasing ecological exposure was significantly associated with reduced distance to wild microbiota (Spearman ρ = −0.48, *p* = 0.003; linear regression slope = −0.031 per exposure level, *p* = 0.0087, R^2^ = 0.19), demonstrating monotonic convergence of microbial community structure toward wild-like states. (**C**) Spearman correlation coefficients (ρ) for top 16 phyla plotted against ecological exposure score (IVC-like < Semi-natural < Eco-tank < Wild). Positive rho values indicate progressive enrichment toward wild conditions; negative values indicate reduction with increasing ecological exposure. *Verrucomicrobiota* (ρ = 0.71, FDR *p* = 2.6 × 10^-5^) and *Desulfobacterota* (ρ = 0.70, FDR *p* = 2.6 × 10^-5^) exhibited strong positive associations, whereas *Bacillota* decreased significantly along the gradient (ρ = −0.63, FDR *p* = 3.1 × 10^-4^). These findings demonstrate coordinated, directional taxonomic restructuring rather than isolated group-specific differences.

### Independent Eco-tanks yield broadly similar gut microbiome configurations

To evaluate the reproducibility of Eco-tank driven gut microbiota preservation, a second identically maintained Eco-tank was established, and microbiomes from the new Eco-tank were compared to those in the first one. Comparison of alpha diversity metrics revealed broadly comparable gut microbiota diversity between wild *Mus* housed in the two independently established Eco-tanks. While Chao1 richness was modestly lower in Eco-tank 2 relative to Eco-tank 1 (Suppl. Figure S3A left), Shannon diversity did not differ significantly between the two groups (*p* = 0.65; Suppl. Figure S3A right), indicating similar community evenness despite minor differences in absolute richness. At the community level (Suppl. Figure S3B), PERMANOVA detected statistically significant differences between tanks using both Bray-Curtis (R^2^ = 0.16, *p* = 0.001) and Jaccard (R^2^ = 0.22, *p* = 0.001) distances. Importantly, homogeneity of dispersion did not differ between tanks (PERMDISP Bray-Curtis *p* = 0.83; Jaccard *p* = 0.155), indicating that observed differences reflect compositional shifts rather than unequal within-group variability. Given that cage effects often contribute 10-30% of total variance in microbiota studies ^62,63^, even among identically maintained IVC cages, our data indicate that independently constructed Eco-tanks reproducibly yield broadly similar gut microbiota profiles with minimal enclosure-specific divergence.

### Eco-tank housing promotes rewilding-like shift of the C57BL/6 gut microbiota

To determine whether sustained ecological exposure can reshape the microbiome of standardized laboratory mice, we evaluated commercially standardized laboratory mice (C57BL/6) housed under IVC, semi-natural, or Eco-tank conditions. Global analysis revealed significant differences in alpha diversity across housing environments (Kruskal-Wallis: Shannon *p* = 0.0006; Chao1 *p* = 8.27 × 10^-5^; **Figure 5A-B**). Pairwise comparisons demonstrated that Eco-tank housing significantly increased both richness and evenness relative to IVC-like cages (Shannon FDR *p* = 0.0019; Chao1 FDR *p* = 0.0015) and semi-natural cages (Shannon FDR *p* = 0.0064; Chao1 FDR *p* = 0.0015) (Suppl. Table S4). In contrast, semi-natural housing did not significantly differ in Shannon diversity compared to IVC-like conditions (FDR *p* = 0.224; Suppl. Table S4), indicating limited restoration of community evenness under partial enrichment.

**Figure 5.**
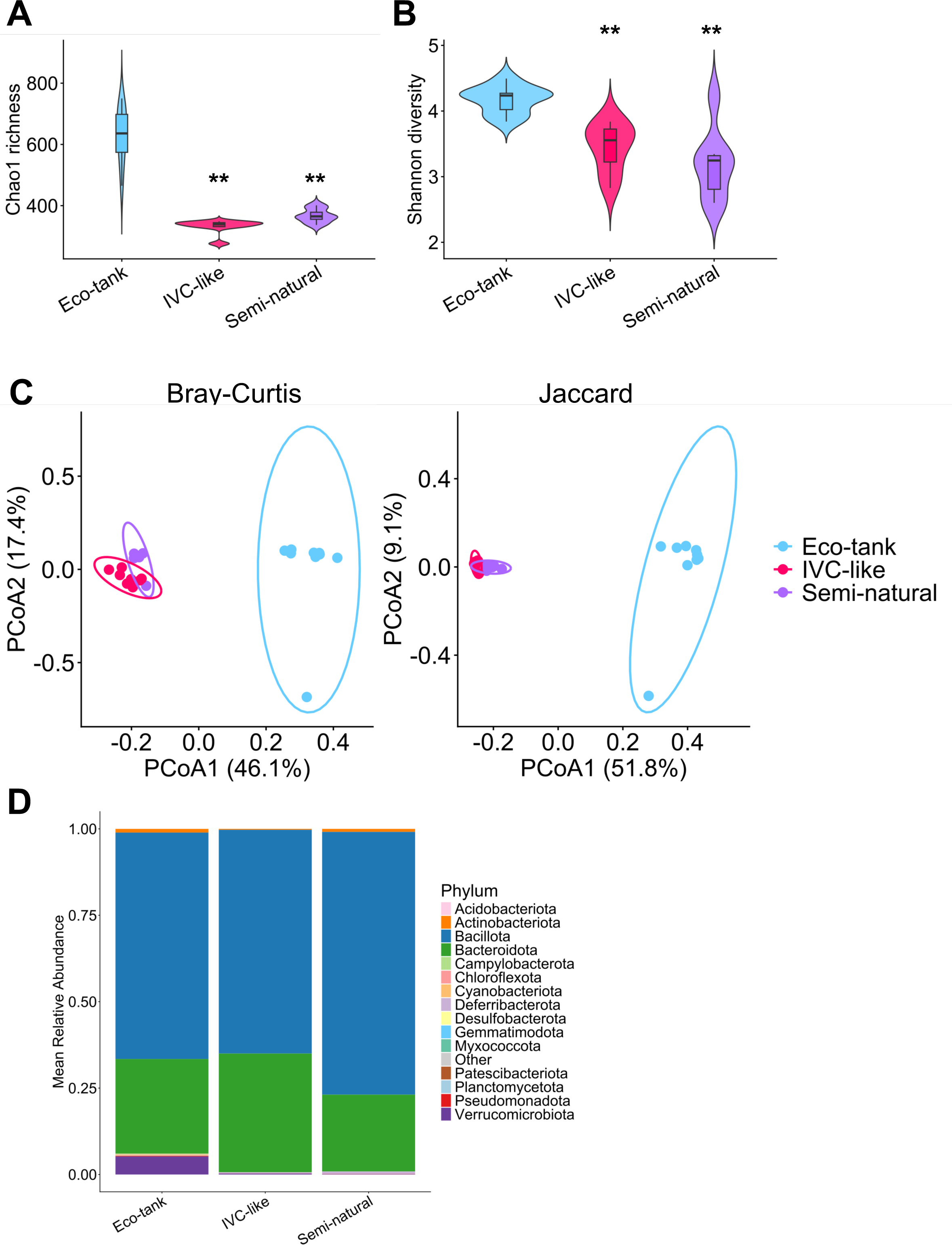
Eco-tank housing induces rewilding-like restructuring of the C57BL/6 gut microbiome. Alpha diversity of gut microbiota across housing conditions was assessed with the Chao1 richness (**A**) and Shannon diversity (**B**) index at ASV levels. Groups include IVC-housing (pink), semi-natural cages (purple), and Eco-tank housing (blue). Violin plots indicate distribution. *Global differences were significant (Kruskal-Wallis, p < 0.001)*. ** *p* < 0.01 compared to Eco-tank based on FDR-adjusted pairwise Wilcoxon tests. (**C**) Principal coordinates analysis (PCoA) of ASV-level community structure based on Bray-Curtis (left; R^2^ = 0.55908, F = 13.314, *p* = 0.001) and Jaccard (right; R^2^ = 0.58496, F = 14.799, *p* = 0.001) dissimilarities. Homogeneity of dispersion was confirmed for Bray-Curtis distances (PERMDISP *p* = 0.288), whereas Jaccard distances exhibited modest but significant differences in dispersion (PERMDISP *p* = 0.005), indicating that presence-absence variation contributed to group separation. Each point represents individual fecal sample, colored by house conditions. Percent variance explained is indicated on the axes. Group differences were tested by PERMANOVA (999 permutations). Ellipses indicate 95% confidence intervals. (**D**) Phylum-level taxonomic composition of gut microbiota across experimental conditions at the indicated time points. Bars represent mean relative abundance across samples within each group. Phylum-level taxonomy was updated to conform with GTDB nomenclature ^84^. Statistical details of this figure are provided in Suppl. Table S4.

Beta diversity analysis further demonstrated clear community-level restructuring (**Figure 5C**). Principal coordinates analysis of Bray-Curtis dissimilarity showed distinct clustering of Eco-tank mice away from IVC-like controls, while semi-natural mice occupying an intermediate but partially overlapping position with IVC housed mice. Jaccard-based analysis yielded similar patterns, indicating that both abundance-weighted and presence-absence community structures were altered under Eco-tank housing (Suppl. Table S4). Homogeneity of dispersion was confirmed for Bray-Curtis distances (PERMDISP *p* = 0.288), whereas Jaccard distances exhibited modest but significant differences in dispersion (PERMDISP *p* = 0.005), indicating that presence-absence variation contributed to group separation.

Consistent with these diversity patterns, phylum-level analysis revealed pronounced compositional differences across housing environments (**Figure 5D**; individual sample data in Suppl. Figure S4A). In IVC-housed C57BL/6 mice, the gut microbiota was dominated by *Bacteroidota* and *Bacillota* with reduced representation of environmentally associated phyla, reflecting an acknowledged laboratory-associated profile. Semi-natural housing partially altered this composition, with increased *Actinobacteriota* (*p* = 0.002) and *Bacillota* (*p* = 0.003) accompanied by a reduction in *Bacteroidota* (*p* = 0.004), but overall taxonomic complexity remained limited. In contrast, Eco-tank housing uniquely promoted the expansion of multiple phyla typically underrepresented or absent in standard laboratory mice (Suppl. Figure S4B), including *Myxococcota* (*p* = 0.0009), *Planctomycetota* (*p* = 0.003), *Pseudomonadota* (*p* = 0.0047), *Verrucomicrobiota* (*p* = 0.0009)*, Acidobacteriota* (*p* = 0.003), *Chloroflexota* (*p* = 0.002), and *Cyanobacteria* (*p* = 0.002). Notably, the Eco-tank was distinguished by the selective enrichment of environmentally responsive and metabolically diverse phyla. Several of these taxa, particularly *Verrucomicrobiota*, *Planctomycetota*, and *Chloroflexota*, were also enriched in wild *Mus* maintained in Eco-tank housing, indicating convergence toward a wild-associated microbial architecture rather than nonspecific diversification.

### Eco-tank housing shifts host-associated microbiota toward an environmental microbial space

To examine the ecological relationship between host-associated gut microbiota and environmental reservoirs, we performed ordination analysis at the ASV level integrating fecal samples from wild-state, IVC-like (wild *Mus* and C57BL/6), and Eco-tank housed mice (wild *Mus* and C57BL/6) with soil-derived microbial communities from the enclosure. Bray-Curtis and Jaccard PCoA revealed clear separation of IVC-like C57BL/6 mice from wild-state animals (**Figure 6A**). In contrast, C57BL/6 mice housed in the Eco-tank displayed a prominent displacement away from the IVC-like cluster and toward the ecological space occupied by wild-state mice. Wild mice maintained in the Eco-tank largely retained proximity to wild-state microbiota profiles. Although soil communities remained distinct from fecal samples, Eco-tank housed mice exhibited reduced visual separation from soil-derived communities relative to IVC-like controls. This shift was evident in both abundance-weighted (Bray-Curtis; **Figure 6A** left) and presence-absence (Jaccard; **Figure 6A** right) metrics, indicating that Eco-tank exposure reshapes both dominant and low-abundance microbial structure.

**Figure 6.**
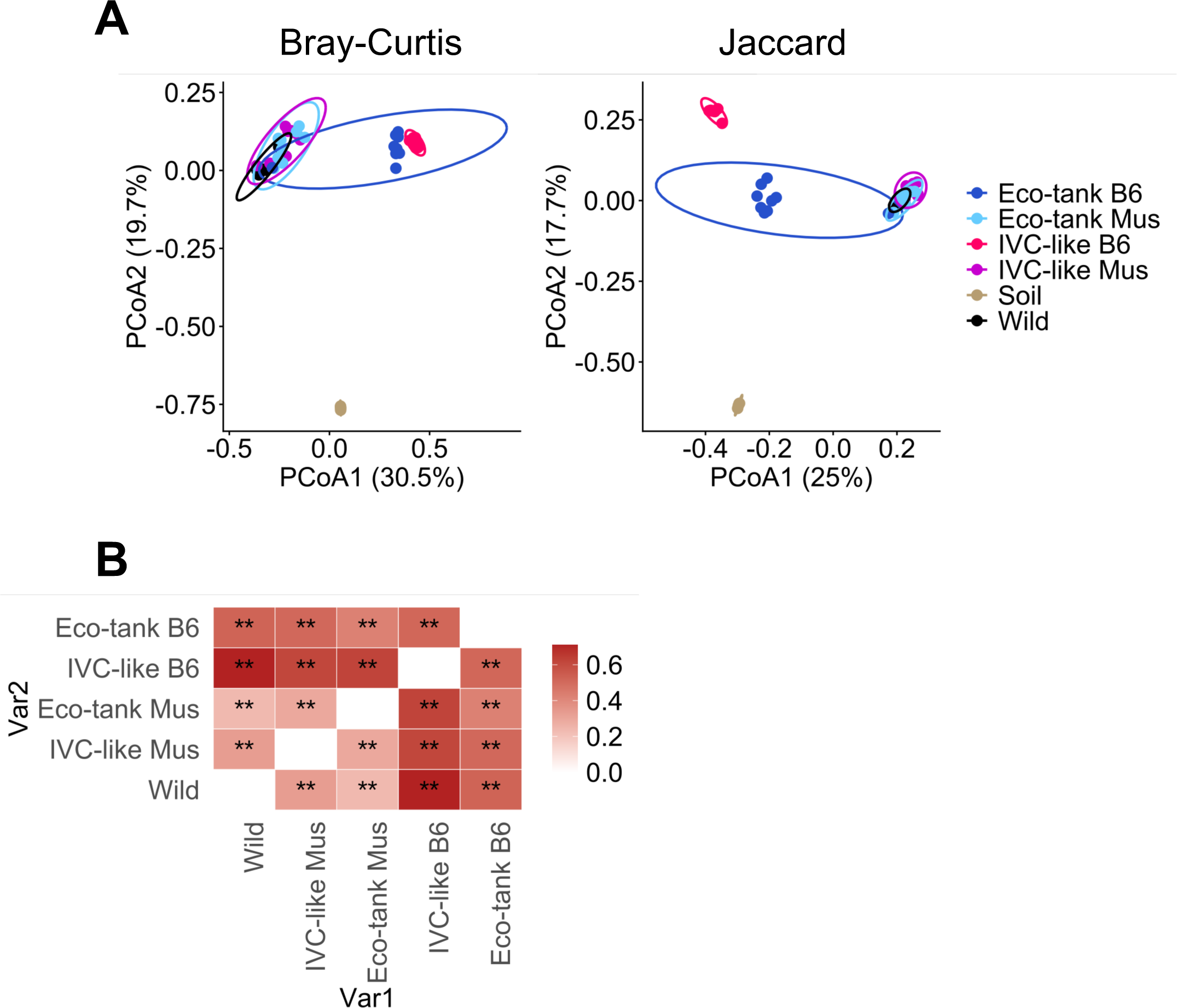
Eco-tank housing repositions the host-associated microbiome toward a wild-associated ecological structure. (**A**) Principal coordinates analysis (PCoA) of ASV-level community structure based on Bray-Curtis (left) and Jaccard (right) dissimilarities. Each point represents an individual sample, colored by housing condition (Wild: Black; IVC-like: Red/Purple; Eco-tank: Blue/Cyan; Soil: Dust Green) and host origin (Wild *Mus*: Black/Purple/Cyan; C57BL/6 or B6: Red/Blue). Ellipses represent 95% confidence intervals. Soil samples were included to visualize ecological proximity and were excluded from statistical testing. Percent variance explained by each axis is indicated in parentheses. (**B**) Pairwise effect size heatmap of housing-dependent microbiota excluding environmental soil samples. Heatmap summarizing pairwise PERMANOVA effect sizes (R^2^) for ASV-level Bray-Curtis (R^2^ = 0.543, *p* = 0.001; PERMDISP *p* = 0.004) across housing conditions. Each tile represents the proportion of variance explained (R^2^) for the comparison between two groups. Warmer colors indicate stronger compositional separation. Statistical significance was determined using permutation-based PERMANOVA with false discovery rate (FDR) correction for multiple comparisons. Asterisks indicate adjusted p-values (**FDR < 0.01). Diagonal entries represent within-group comparisons and are shown as zero. Statistical details of this figure are provided in Suppl. Table S5.

To quantify the contribution of housing condition to gut microbial community structure, we performed PERMANOVA analyses excluding environmental soil samples. Housing condition explained a substantial proportion of variance in community composition based on both Bray-Curtis (R^2^ = 0.543, *p* = 0.001; Suppl. Table S5) and Jaccard distance (R^2^ = 0.555, *p* = 0.001; Suppl. Table S5), indicating that environmental context is a dominant determinant of microbiota architecture across host origins. Pairwise PERMANOVA comparisons revealed a clear hierarchy of divergence (Bray-Curtis; **Figure 6B**; Suppl. Table S5). IVC-like B6 mice exhibited the greatest compositional separation from wild-state and Eco-tank groups (R^2^ up to 0.711), consistent with pronounced ecological constraint under standard laboratory housing. In contrast, divergence between wild-state and Eco-tank housed wild *Mus* was substantially reduced (R^2^ = 0.132), indicating partial preservation of wild-like microbiome structure under Eco-tank conditions (Jaccard in Suppl. Table S5). Assessment of homogeneity of multivariate dispersion demonstrated modest but significant differences among groups for both Bray-Curtis (F = 4.61, *p* = 0.004; Suppl. Table S5) and Jaccard distance (F = 6.31, *p* = 0.001; Suppl. Table S5). Dispersion differences were primarily driven by IVC-like B6 mice, which exhibited reduced within-group variability relative to Eco-tank and wild-state mice. Despite these differences in dispersion, the large effect sizes observed in PERMANOVA analyses support robust centroid separation and substantive ecological restructuring across housing conditions.

We further showed that the Eco-tank reshaped the microbiota in a genotype-dependent manner using within-genotype analyses (Suppl. Table S5). Factorial PERMANOVA revealed a significant Genotype × Housing interaction (excluded wild state; R^2^ = 0.091, *p* = 0.001; Suppl. Table S5). Eco-tank housing explained 51.3% of microbiota change in C57BL/6 mice and 29.3% in wild-derived *Mus* (both PERMANOVA, *p* = 0.001). Dispersion did not differ significantly among groups (PERMDISP *p* = 0.067). C57BL/6 exhibited greater ecological plasticity in response to Eco-tank exposure, with the magnitude of restructuring was nearly two-fold stronger than in wild-derived *Mus*.

### Functional association analysis reveals coordinated pathway shifts in wild *Mus* upon housing condition changes

To determine whether Eco-tank housing mitigates functional erosion observed under IVC conditions, we examined predicted KEGG pathway enrichment across housing states using TAX4FUN2. Eco-tank replicate cohorts (Eco-tank 1 and Eco-tank 2), along with the IVC cohorts at 14 days and at 52 days were analyzed separately at the ASV level to confirm consistency. As predicted functional profiles did not differ significantly between Eco-tank replicates (PERMANOVA, F = 1.84, R^2^ = 0.098, *p* = 0.16; PERMDISP, *p* = 0.97), as well as different timepoints of IVC housed mice data (PERMANOVA, F = 0.36, R^2^ = 0.029, *p* = 0.702; PERMDISP, *p* = 0.99), cohorts were combined for downstream functional analysis to maximize statistical power (Suppl. Figure S5). Principal component analysis (PCA) of predicted KEGG Ortholog profiles revealed clear clustering of samples according to housing condition (**Figure 7A**). Housing conditions explained 35.1% of the variance in functional composition across groups (PERMANOVA, F = 9.75, R^2^ = 0.351, *p* = 0.001; Suppl. Table S6). Microbial communities from wild-state *Mus* occupied a distinct region of functional space compared to IVC-housed mice (F = 21.80, R^2^ = 0.548, FDR = 0.0015; Suppl. Table S6), indicating substantial differences in predicted metabolic capacity. Eco-tank mice displayed an intermediate functional profile, partially overlapping with both wild-state (F = 2.56, R^2^ = 0.100, FDR = 0.051) and IVC groups (F = 11.36, R^2^ = 0.268, FDR = 0.0015), suggesting partial retention or reacquisition of functional traits characteristic of the wild microbiome. Homogeneity of dispersion was largely comparable across groups (PERMDISP, *p* = 0.059), although a modest difference was detected between IVC and Eco-tank samples (PERMDISP, *p* = 0.042).

**Figure 7.**
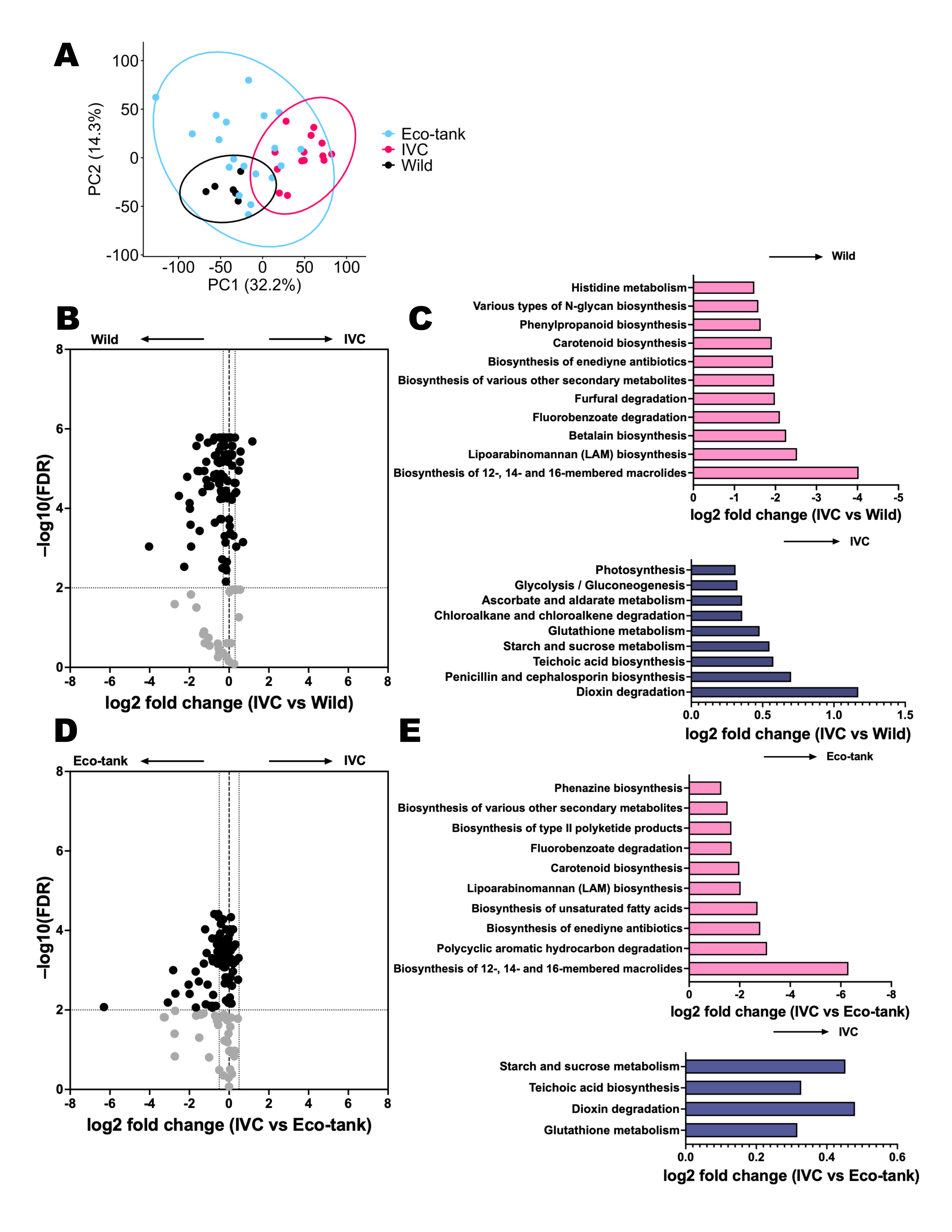
Eco-tank housing preserves wild-like predicted microbial functional profiles in *wild Mus*. (**A**) Principal component analysis (PCA) of predicted functional profiles generated using TAX4FUN2 from ASV-level data. Functional abundances (KEGG Ortholog level) were normalized to relative abundance and scaled prior to analysis. Each point represents one fecal sample. Percent variance explained by each principal component is indicated on the axes. Group differences in overall functional composition were assessed using Bray-Curtis dissimilarity and PERMANOVA (999 permutations) (PERMANOVA, F = 9.75, R^2^ = 0.351, *p* = 0.001; PERMDISP, *p* = 0.059). Ellipses indicate 95% confidence intervals. Statistical details of this panel are provided in Suppl. Table S6. (**B**) Volcano plots show differential predicted KEGG functional pathways between wild-state and after housing in IVC cages for 52 days. Each point represents a KEGG pathway predicted by Tax4Fun. The x-axis shows log2 fold change (IVC / Wild), and the y-axis shows -log10(FDR). Pathways enriched in wild state are shown on the left, while pathways enriched under IVC conditions are shown on the right. (**C**) Bar plots highlighting representative pathways significantly enriched in Wild (top) or IVC-like cage (bottom) conditions. All pathways shown met an FDR threshold of <0.01. (**D**) Volcano plot showing differential predicted KEGG functional pathways between Eco-tank and IVC cages. Each point represents a KEGG pathway predicted by Tax4Fun. The x-axis shows log2 fold change (IVC / Eco-tank), and the y-axis shows -log10(FDR). Pathways enriched in Eco-tank mice are shown on the left, while pathways enriched under IVC conditions are shown on the right. (**E**) Bar plots highlighting representative pathways significantly enriched in Eco-tank (top) or IVC-like cage (bottom) conditions. All pathways shown met an FDR threshold of <0.01.

To identify predicted microbial functions most strongly associated with the transition from a wild environment to IVC housing, we performed functional association analysis using a global test framework. This approach identified multiple KEGG pathways whose predicted abundances were significantly associated with housing conditions. Differential analysis of these functionally associated pathways revealed pronounced and directional shifts following IVC housing (**Figure 7B**). Numerous pathways showed significantly higher predicted abundance in the wild state (**Figure 7C** top; log2 fold < 0, FDR < 0.05), including pathways involved in secondary metabolite biosynthesis, amino acid metabolism, complex glycan biosynthesis, and xenobiotic degradation. In contrast, pathways enriched after IVC housing were primarily associated with central carbon metabolism and redox-related processes, including glycolysis/gluconeogenesis, starch and sucrose metabolism, and glutathione metabolism (**Figure 7C** bottom; log2 fold > 0, FDR < 0.05).

To determine whether Eco-tank housing mitigates the functional losses associated with IVC conditions, we compared predicted microbial functional profiles between Eco-tank and IVC mice using functional association analysis (**Figure 7D**). Multiple pathways involved in secondary metabolite biosynthesis, xenobiotic degradation, lipid metabolism, and complex antimicrobial compound production were significantly enriched in Eco-tank mice relative to IVC mice (**Figure 7E** top; log2 fold < 0, FDR < 0.05). In contrast, IVC mice exhibited higher predicted abundance of pathways associated with central carbohydrate metabolism and redox homeostasis, including starch and sucrose metabolism, teichoic acid biosynthesis, and glutathione metabolism (**Figure 7E** bottom; log2 fold > 0, FDR < 0.05). The magnitude and directionality of these changes indicate that Eco-tank housing retains a broader and more environmentally representative functional repertoire than IVC housing, while still differing from fully wild microbiomes.

### Eco-tank housing preserves microbial pathways involved in different metabolic states but incompletely maintains vitamin and antioxidant pathways

We then focused on predicted enrichment of SCFA production (propanoate, butanoate metabolism), mucin- and glycan-targeting pathways (amino sugar and nucleotide sugar metabolism, other glycan degradation), and tryptophan and glutathione metabolism, all of which are known to support mucosal IgA and IL-17/IL-22 mediated barrier function ^64–67^. Under IVC housing, wild-derived microbiota exhibited marked depletion of core energy metabolism pathways, including fatty acid degradation, glyoxylate and dicarboxylate metabolism, and the citrate (TCA) cycle (Suppl. Figure S6A). These pathways are central to microbial carbon flux and short-chain fatty acid (SCFA) production ^66,68^. In contrast, Eco-tank housing preserved multiple components of this metabolic network (Suppl. Figure S6B), suggesting sustained microbial energy metabolism under semi-naturalistic conditions.

A similar outcome was observed for amino acid and tryptophan metabolism. IVC housing resulted in significant depletion of histidine metabolism, branched-chain amino acid biosynthesis, lysine degradation, and tryptophan metabolism (Suppl. Figure S6C). Eco-tank housing mitigated this loss, maintaining enrichment of several amino acid pathways (Suppl. Figure S6D), including tryptophan metabolism ^69^, which is closely linked to mucosal immune modulation through aryl hydrocarbon receptor (AhR) signaling ^70^. Functional categories related to microbial structural components and PAMP biosynthesis further distinguished housing states. Wild-state microbiota was enriched for lipopolysaccharide (LPS) and exopolysaccharide biosynthesis, as well as glycosaminoglycan degradation and galactose metabolism (Suppl. Figure S6E and G). These features were markedly diminished under IVC conditions. Eco-tank housing partially restored LPS biosynthesis and glycosaminoglycan degradation pathways relative to IVC mice (Suppl. Figure S6F and H), indicating retention of structural microbial features associated with mucosal immune engagement ^71^.

Environmental and xenobiotic metabolism pathways also differed by housing condition. Fluorobenzoate degradation was enriched in wild-state microbiota but reduced under IVC conditions (Suppl. Figure S6I). Eco-tank housing preserved this pathway and additionally enriched benzoate degradation relative to IVC-like mice (Suppl. Figure S6J), suggesting sustained environmental metabolic capacity without complete recapitulation of the wild-state functional landscape. In contrast to metabolic and structural pathways, vitamin and antioxidant pathways showed incomplete preservation. Wild-state microbiota was enriched for vitamin B6 and riboflavin metabolism, both of which were markedly reduced under IVC housing (Suppl. Figure S6K). Eco-tank housing did not fully restore these pathways (Suppl. Figure S6L), indicating selective rather than global preservation of wild-associated functional traits.

### Environmental enrichment reshapes predicted microbial functional profiles in C57BL/6 mice

To determine whether environmental complexity alters microbial functional potential in a standard laboratory strain, we inferred predicted microbial functions in C57BL/6 mice housed under IVC, semi-natural, or Eco-tank conditions. Principal component analysis of predicted KEGG functional profiles revealed clear separation of samples by housing condition (F = 13.314, R^2^ = 0.55908, *p* = 0.001; PERMDISP *p* = 0.263; **Figure 8A**). IVC-housed B6 mice clustered tightly, indicating constrained functional diversity under standard laboratory conditions. In contrast, Eco-tank mice occupied a distinct region of ordination space, reflecting increased heterogeneity in predicted microbial metabolic capacity (Suppl. Table S7).

**Figure 8.**
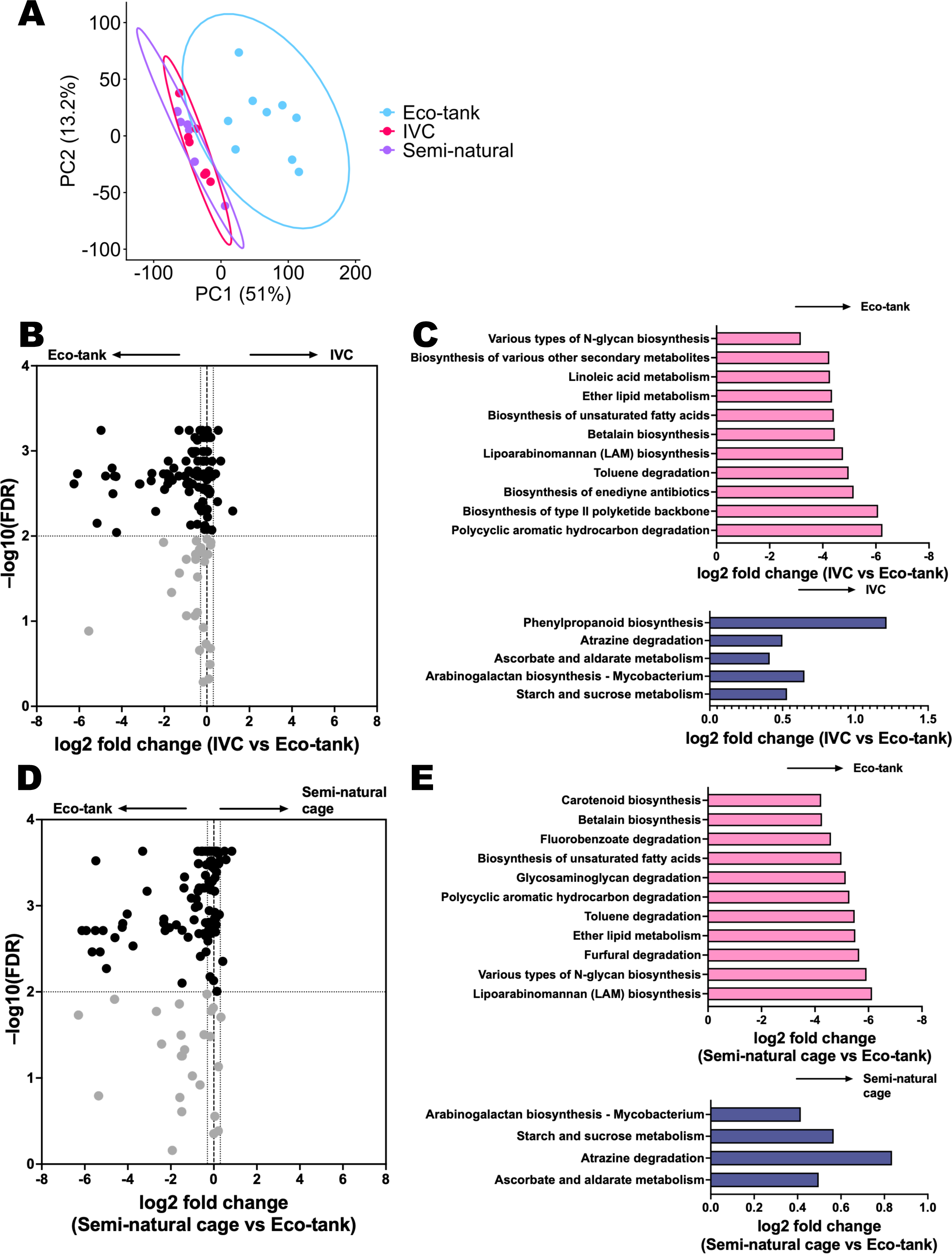
Eco-tank housing induces coordinated functional reprogramming of the gut microbiome in C57BL/6 mice. (**A**) Principal component analysis (PCA) of predicted functional profiles generated using TAX4FUN2 from ASV-level data. Functional abundances (KEGG Ortholog level) were normalized to relative abundance and scaled prior to analysis. Each point represents one fecal sample. Percent variance explained by each principal component is indicated on the axes. Group differences in overall functional composition were assessed using Bray-Curtis dissimilarity and PERMANOVA (999 permutations) (R^2^ = 0.55908, F = 13.314, *p*= 0.001; PERMDISP *p* = 0.263). Ellipses indicate 95% confidence intervals. Statistical details of this panel are provided in Suppl. Table S7. (**B**) Volcano plot showing differential predicted KEGG functional pathways between Eco-tank and IVC cages. Each point represents a KEGG pathway predicted by Tax4Fun. The x-axis shows log2 fold change (IVC / Eco-tank), and the y-axis shows -log10(FDR). Pathways enriched in Eco-tank mice are shown on the left, while pathways enriched under IVC conditions are shown on the right. (**C**) Bar plots highlighting representative pathways significantly enriched in Eco-tank (top) or IVC-like cage (bottom) conditions. All pathways shown met an FDR threshold of <0.01. (**D**) Volcano plot showing differential predicted KEGG pathways between B6 mice housed in semi-natural cages and Eco-tank environments. Pathway-level abundances inferred using Tax4Fun were used to calculate log2 fold changes (Semi-natural cage / Eco-tank). The x-axis shows log2 fold change, and the y-axis shows -log10(FDR). Each point represents a KEGG pathway; black points indicate pathways meeting the significance threshold (FDR-adjusted). Pathways enriched in Eco-tank mice are shown on the left, while pathways enriched under semi-natural conditions are shown on the right. (**E**) Bar plots highlight representative pathways significantly enriched in Eco-tank (top) or semi-natural cage (bottom) conditions. All pathways shown met an FDR threshold of <0.01.

We then compared predicted microbial pathways between IVC- and Eco-tank housed C57BL/6 mice using functional association analysis. Volcano plot visualization revealed widespread and directional changes in predicted KEGG pathways following transfer from IVC housing to Eco-tank conditions (**Figure 8B**). Multiple pathways were significantly enriched in Eco-tank housed C57BL/6 mice, including pathways involved in secondary metabolite biosynthesis, complex lipid metabolism, glycan biosynthesis, and fatty acid biosynthesis (**Figure 8C** top; log2 fold < 0, FDR < 0.05), indicating expansion of microbial metabolic breadth under Eco-tank housing. Eco-tank housed mice also showed significantly enriched pathways relative to semi-natural cage mice (**Figure 8D**), including complex lipid metabolism, glycan biosynthesis, fatty acid biosynthesis, as well as polycyclic aromatic hydrocarbon and toluene degradation (**Figure 8E** top; log2 fold < 0, FDR < 0.05). In contrast, IVC-housed C57BL/6 mice were enriched for a smaller subset of pathways primarily associated with central carbohydrate metabolism and select biosynthetic functions compared to both Eco-tank housing (**Figure 8C** bottom; log2 fold > 0, FDR < 0.05) and semi-natural cages (**Figure 8E** bottom; log2 fold > 0, FDR < 0.05), including starch and sucrose metabolism, arabinogalactan biosynthesis, and atrazine degradation. These differences indicated that Eco-tank and semi-natural environments promote distinct functional configurations of the gut microbiome rather than representing identical forms of environmental enrichment.

### Eco-tank housing yields vaccine outcomes consistent with environmentally conditioned C57BL/6 mouse immunity

To test whether housing-dependent microbiome differences impact vaccine efficacy, C57BL/6 mice housed in semi-natural cages or Eco-tank environments were immunized intranasally with L-PaF formulated in BECC/ME (BRAVE.0) and subsequently challenged with *P. aeruginosa* (Pa). In semi-natural caged mice, vaccination resulted in a pronounced reduction in lung bacterial burden compared to naïve controls (**Figure 9A**). Vaccinated animals exhibited near-complete clearance of Pa, whereas naïve mice displayed high and variable lung CFU counts. In Eco-tank housed mice, vaccination also significantly reduced pulmonary bacterial burden relative to naïve controls (**Figure 9B**). Although naïve Eco-tank mice exhibited lower baseline CFU counts compared to naïve semi-natural caged mice, vaccinated Eco-tank mice showed a further and statistically significant reduction in bacterial load. These results indicate that Eco-tank housing supports effective vaccine-induced protection against Pa lung infection in C57BL/6 mice and suggest that environmental housing conditions influence both baseline susceptibility and vaccine responsiveness.

**Figure 9.**
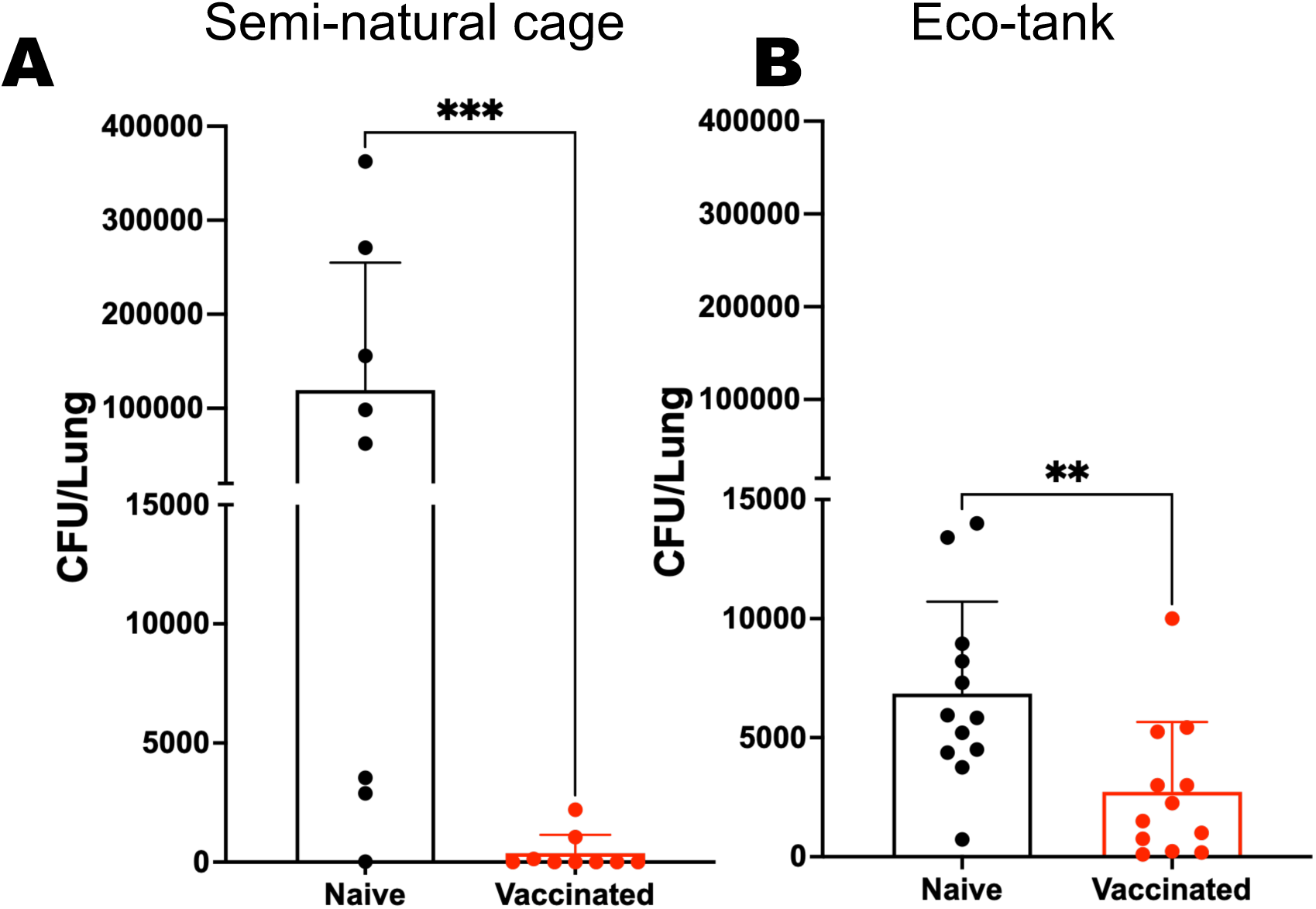
Eco-tank housing represents a balanced model against pulmonary *Pseudomonas aeruginosa* (Pa) infection in C57BL/6 mice. C57BL/6 mice housed in semi-natural cages (**A**; n = 8-9/group) or Eco-tank environments (**B**; n = 12/group) were immunized intranasally with BRAVE.0 or received PBS as a control. Mice were challenged intranasally with 4 × 10^7^ CFU of Pa, and lung bacterial burden (CFU) was quantified 24 hours post-infection. Each point represents a sample from an individual mouse; bars indicate mean ± SEM. Statistical significance was assessed between naïve and vaccinated groups by Mann-Whitney test (\*\**p* < 0.01, \*\*\**p* < 0.001).

Notably, the level of protection observed under semi-natural housing was comparable to that previously reported in IVC-housed mice ^50–53^, whereas Eco-tank housed mice displayed a distinct outcome profile. This pattern parallels the clustering observed in predicted functional microbiota profiles. Functional PCA demonstrated close grouping of IVC-like and semi-natural samples, while Eco-tank samples formed a separate cluster (**Figure 8A**). These findings suggest that similarity in predicted microbial functional architecture may be associated with similarity in vaccine-mediated protection phenotypes.

## Discussion

The modern laboratory environment constitutes a profound ecological bottleneck for host-associated microbial communities. While specific pathogen free rearing followed by on site IVC housing has enabled experimental reproducibility in immunology, it has simultaneously constrained microbial diversity and functional breadth ^72,73^. Our findings demonstrate that housing conditions are not passive background variables but dominant ecological forces that restructure gut microbiome composition, reshape predicted functional capacity, and alter host immune outcomes. Consistent with this view, transition from wild environments to IVC housing resulted in rapid contraction of microbial diversity and marked compositional remodeling. These taxonomic shifts were accompanied by coordinated reductions in predicted pathways related to central carbon metabolism, short-chain fatty acid (SCFA) production, amino acid metabolism, and structural microbial components such as lipopolysaccharide and exopolysaccharide biosynthesis. Although derived from 16S-based inference, the consistency of these directional changes suggests that traditional laboratory housing compresses both the structural and metabolic landscape of the microbiome.

In contrast, the Eco-tank system disrupts this constrained state. In wild-derived mice, Eco-tank housing was found to stabilize microbial diversity and preserve elements of wild-associated taxonomic architecture during extended captivity. In standard C57BL/6 mice, Eco-tank also drove a rewilding-like restructuring of the microbiome, increasing richness and evenness while expanding environmentally associated phyla rarely observed under IVC conditions. Moreover, this shift was substantially stronger than that observed in wild-derived *Mus*, indicating a greater ecological plasticity for the laboratory mouse microbiota. Notably, partial enrichment through soil or diet alone was insufficient; only sustained ecological complexity produced robust and reproducible remodeling, with increasing ecological complexity associated with reduced distance to wild communities. Consistent with this gradient, several phyla - including *Desulfobacterota* ^74^, *Verrucomicrobiota*^75^, and *Actinobacteriota* ^76^ - displayed directional enrichment across housing conditions. These taxa were commonly associated with anaerobic environmental metabolism, mucosal interface interactions, and secondary metabolite production, suggesting that sustained ecological complexity re-establishes metabolic niches that remain inaccessible under partial enrichment conditions. Consistent with these taxonomic shifts, structural microbial features also shifted across housing conditions. Wild-associated communities were enriched in Gram-negative associated biosynthetic pathways, whereas IVC housing favored signatures consistent with Gram-positive dominated laboratory microbiota. Eco-tank housing partially restored Gram-negative associated features while retaining elements of IVC architecture, generating a diversified microbial landscape with potential to modulate innate immune conditioning ^77^. These observations suggest that ecological complexity influences not only microbial metabolism but also the spectrum of microbial-associated molecular patterns interfacing with host immunity ^77,78^.

Notably, Eco-tank microbiomes did not simply replicate the wild state. Instead, they established a distinct ecological configuration characterized by expansion of lipid metabolism, glycan utilization, aromatic compound degradation, and select secondary metabolite pathways. Certain pathways were uniquely enriched under Eco-tank conditions, indicating that controlled environmental complexity imposes distinct selective pressures rather than simply restoring the pre-capture wild state. Even in genetically identical C57BL/6 mice, Eco-tank housing substantially expanded the predicted functional repertoire of the gut microbiome. Eco-tank housing enriched predicted pathways associated with SCFA production ^67^, amino acid metabolism ^79^, and degradation of environmentally derived compounds ^80^, features that were more characteristic of human-associated microbiomes than tightly constrained IVC communities. Although these functional predictions require direct metabolomic validation, their enrichment aligns with the enhanced baseline resistance to pulmonary *P. aeruginosa* observed in Eco-tank housed mice. Importantly, vaccine-induced protection remained intact though it resembled what one might expect in a genetically diverse outbred population, indicating that restoration of environmental microbial signals does not compromise overall adaptive responsiveness. Instead, ecological conditioning broadened baseline immune competence while preserving vaccine efficacy.

Interestingly, vaccine protection phenotypes paralleled microbiome functional organization across housing conditions. Semi-natural housing recapitulated the BRAVE.0 vaccine protection observed under IVC conditions and clustered closely with IVC microbiota at the predicted functional level, whereas Eco-tank microbiota adopted a distinct functional configuration accompanied by divergence in early antibacterial protection. Although causality cannot be established in this study, the concordance between microbial functional clustering and protection outcomes suggests that baseline microbial metabolic tone may influence the kinetics or magnitude of vaccine-elicited responses. Given that L-PaF-mediated protection involves Th17, Th1/Th2, and TCR-associated signaling pathways ^50,53^, which are immune axes known to be modulated by microbiota-derived signals ^5,65^, environmentally structured microbiome may shape vaccine-induced immunity in ways not captured under standard laboratory housing.

From a translational perspective, the Eco-tank model offers a deliberate compromise between ecological realism and experimental control. Unlike wild or fully rewilded models, which introduce genetic and environmental variability^16,19–23^, the Eco-tank is constructed from defined components and operated under pathogen-monitored conditions. Although tank identity contributed modestly to community variance, which was within the range commonly observed for enclosure effects in animal studies ^63,81^, the overall system remained reproducible. This balance of biosecurity, scalability, and ecological complexity positions the Eco-tank as a practical intermediate platform for studying microbiome-driven immune regulation under conditions more reflective of real-world microbial exposure. Importantly, intestinal microbiome composition is increasingly recognized as a determinant of inter-individual variability in vaccine-induced immunity ^82^. By broadening microbial functional potential without compromising adaptive responsiveness, Eco-tank housing may provide a more translationally relevant baseline for preclinical vaccine evaluation.

Yet few limitations warrant consideration. Functional profiles were inferred from 16S rRNA gene data and therefore represent predicted metabolic potential rather than measured activity. Direct metagenomic and metabolomic analyses will be required to validate pathway function and identify specific metabolites mediating immune effects. In addition, mechanistic dissection of the causal links between microbial restructuring and enhanced baseline immunity remains an important future direction. Nonetheless, the directional consistency of these shifts supports the broader principle that ecological context, even under controlled laboratory conditions, is a dominant force shaping microbiome structure and functional potential ^83^.

In summary, our findings support a model in which laboratory housing imposes ecological constraint on the gut microbiome, compressing diversity and functional breadth into a simplified, laboratory-adapted configuration. Eco-tank housing selectively reverses key aspects of this constraint, establishing an intermediate yet functionally expanded ecological state that broadens host-microbe interactions while maintaining experimental tractability. Incorporating environmental context as a deliberate experimental variable may help realign preclinical immunology with the ecological realities that shape immune function in natural settings.

## Author contributions

T.L. was responsible for conceptualization, data acquisition, curation, data analysis, method development, data visualization and writing of the initial manuscript draft; W.L.P. was responsible for acquiring financial support, project administration, editing of the original manuscript draft and overall supervision; Z.K.D. was responsible for the collection of specimens, protein purification, ME nanoemulsion preparation, and animal housing and method development; A.C.E. was responsible for data generation and curation, and critical reading and editing of the manuscript; W.D.P was responsible for final manuscript editing and writing.

## Acknowledgements.

This work was funded by NIAID grants R01AI138970 to WLP. We thank the MU Veterinarians and support staff of the MU Office of Animal Resource for their guidance, helpful discussions, and flexibility in supporting the development and implementation of the eco-tank housing system. We also thank Dr. Robert K. Ernst in the University of Maryland for the resource of BECC438s.

## Conflict

WLP and WDP are affiliated with/cofounders of Hafion, Inc. The remaining authors declare that the research was conducted in the absence of any commercial or financial relationships that could be construed as a potential conflict of interest.

## Data Availability Statement

The original contributions presented in the study are included in the article/supplementary material, further inquiries can be directed to the corresponding authors. The datasets generated and/or analyzed during the current study will be made available by the corresponding author upon reasonable request after the manuscript has been accepted for publication.

