## Supplemental Tables & Figures for "Eco-tank Housing Maintains Wild-Type Microbiota and Rewilds the Laboratory Mouse Gut Microbiome to Restore Natural Immune Tone"

**Suppl. Table S1.** Abbreviations

| **Abbreviation** | **Full Term** |
| --- | --- |
| ASV | Amplicon Sequence Variant |
| IVC | Individually ventilated cage |
| TSS | Total Sum Scaling |
| KO | KEGG Ortholog |
| KEGG | Kyoto Encyclopedia of Genes and Genomes |
| PCoA | Principal Coordinates Analysis |
| PCA | Principal Component Analysis |
| PERMANOVA | Permutational Multivariate Analysis of Variance |
| PERMDISP | Permutational Analysis of Multivariate Dispersions |
| FDR | False Discovery Rate |
| Chao1 | Chao1 Richness Estimator |
| R^2^ | Coefficient of Determination |
| Rho or ρ | Spearman correlation coefficients |
| β | Slope of linear modeling |
| Lwr | Lower Confidence Interval |
| Upr | Upper Confidence Interval |
| Wild *Mus* | wild *Mus musculus* |
| Wild | wild *Mus musculus* (*Mus*) at the time of capture |
| SEM | Standard Error of the Mean |
| IQR | Interquartile Range |
| AhR | Aryl Hydrocarbon Receptor |
| SCFA | Short-Chain Fatty Acid |
| 16S rRNA | 16S Ribosomal RNA Gene |
| Bray-Curtis | Bray-Curtis Dissimilarity |
| Jaccard | Jaccard Distance Index |
| L-PaF | *Pseudomonas aeruginosa* vaccine Formulation with LTA1 |
| BECC | Bacterial Enzymatic Combinatorial Chemistry |
| ME | squalene-based MedImmune Emulsion |
| Pa | *Pseudomonas aeruginosa* |
| LTA1 | a mucosal adjuvant derived from the double-mutant labile toxin (dmLT) |

**Suppl. Table S2. Pathogen screening results for wild mice using the IDEXX Mouse Global PCR panel. The complete diagnostic report is available from the authors upon request.**

| **Agent** | **Result** |
| --- | --- |
| *Salmonella spp.* | Negative |
| *Campylobacter spp.* | Negative |
| *Pseudomonas aeruginosa* | Negative |
| *Staphylococcus aureus* | Negative |
| *Streptobacillus moniliformis* | Negative |
| *Helicobacter hepaticus* | Positive |
| *Helicobacter ganmani* | Positive |
| *Rodentibacter pneumotropicus* | Positive |
| MPV | Positive |

**Suppl. Table S3.** Statistics for Figure 3. All analyses were performed at the ASV level.

A. Alpha Diversity (Kruskal-Wallis test followed by pairwise Wilcoxon rank-sum tests with FDR correction.)

Shannon Diversity (FDR-adjusted p values)

| **Comparison** | **FDR-adjusted p** |
| --- | --- |
| Semi-natural vs Eco-tank | 0.1177 |
| Soil bedding vs Eco-tank | 0.0021 |
| IVC-like vs Eco-tank | 0.0029 |
| Naturalistic diet vs Eco-tank | 0.0091 |
| Wild vs Eco-tank | 0.7286 |
| Soil bedding vs Semi-natural | 0.0029 |
| IVC-like vs Semi-natural | 0.245 |
| Naturalistic diet vs Semi-natural | 0.9698 |
| Wild vs Semi-natural | 0.1177 |
| IVC-like vs Soil bedding | 0.0627 |
| Naturalistic diet vs Soil bedding | 0.0021 |
| Wild vs Soil bedding | 0.0029 |
| Naturalistic diet vs IVC-like | 0.118 |
| Wild vs IVC-like | 0.003 |
| Wild vs Naturalistic diet | 0.0029 |

Chao1 Richness (FDR-adjusted p values)

| **Comparison** | **FDR-adjusted p** |
| --- | --- |
| Semi-natural vs Eco-tank | 0.0026 |
| Soil bedding vs Eco-tank | 0.0014 |
| IVC-like vs Eco-tank | 0.0014 |
| Naturalistic diet vs Eco-tank | 0.0014 |
| Wild vs Eco-tank | 0.953 |
| Soil bedding vs Semi-natural | 0.006 |
| IVC-like vs Semi-natural | 0.0615 |
| Naturalistic diet vs Semi-natural | 0.1397 |
| Wild vs Semi-natural | 0.0026 |
| IVC-like vs Soil bedding | 0.0616 |
| Naturalistic diet vs Soil bedding | 0.0069 |
| Wild vs Soil bedding | 0.0026 |
| Naturalistic diet vs IVC-like | 0.293 |
| Wild vs IVC-like | 0.003 |
| Wild vs Naturalistic diet | 0.0026 |

B. Beta Diversity - Bray-Curtis (PERMANOVA, 999 permutations)

Overall Model

| **Source** | **Df** | **Sum of Squares** | **R^2^** | **F** | **p** |
| --- | --- | --- | --- | --- | --- |
| Model | 5 | 4.3531 | 0.4381 | 7.642 | 0.001 |
| Residual | 49 | 5.5823 | 0.5619 |  |  |
| Total | 54 | 9.9353 | 1 |  |  |

Pairwise PERMANOVA (FDR-adjusted)

| **Group 1** | **Group 2** | **F** | **R^2^** | **p** | **FDR** |
| --- | --- | --- | --- | --- | --- |
| Wild | Soil bedding | 12.73 | 0.476 | 0.001 | 0.001 |
| Wild | Naturalistic diet | 7.18 | 0.339 | 0.001 | 0.001 |
| Wild | Semi-natural | 5.15 | 0.269 | 0.001 | 0.001 |
| Wild | IVC-like | 6.95 | 0.332 | 0.001 | 0.001 |
| Wild | Eco-tank | 4.03 | 0.237 | 0.001 | 0.001 |
| Soil bedding | Naturalistic diet | 16.66 | 0.481 | 0.001 | 0.001 |
| Soil bedding | Semi-natural | 12.5 | 0.41 | 0.001 | 0.001 |
| Soil bedding | IVC-like | 3.12 | 0.148 | 0.002 | 0.002 |
| Soil bedding | Eco-tank | 11.22 | 0.398 | 0.001 | 0.001 |
| Naturalistic diet | Semi-natural | 4.36 | 0.195 | 0.001 | 0.001 |
| Naturalistic diet | IVC-like | 7.54 | 0.295 | 0.001 | 0.001 |
| Naturalistic diet | Eco-tank | 7.63 | 0.31 | 0.001 | 0.001 |
| Semi-natural | IVC-like | 6.15 | 0.255 | 0.001 | 0.001 |
| Semi-natural | Eco-tank | 6.29 | 0.27 | 0.001 | 0.001 |
| IVC-like | Eco-tank | 7.05 | 0.293 | 0.001 | 0.001 |

C. Homogeneity of Dispersion (PERMDISP)

No significant differences in dispersion were detected between groups (all adjusted p > 0.28).

D. Beta Diversity -Jaccard (PERMANOVA, 999 permutations)

| **Source** | **Df** | **Sum of Squares** | **R^2^** | **F** | **p** |
| --- | --- | --- | --- | --- | --- |
| Model | 5 | 2.9673 | 0.3792 | 5.987 | 0.001 |
| Residual | 49 | 4.857 | 0.6208 |  |  |
| Total | 54 | 7.8243 | 1 |  |  |

**Suppl. Table S4** Statistical analysis for Figure 5. All analyses were performed at the ASV level.

A. Alpha Diversity (Kruskal-Wallis with FDR-adjusted pairwise Wilcoxon tests)

Shannon Diversity (FDR-adjusted p values)

| **Comparison** | **FDR-adjusted p** |
| --- | --- |
| Semi-natural vs Eco-tank | 0.0064 |
| IVC-like vs Eco-tank | 0.0019 |
| IVC-like vs Semi-natural | 0.2243 |

Chao1 Richness (FDR-adjusted p values)

| **Comparison** | **FDR-adjusted p** |
| --- | --- |
| **Semi-natural vs Eco-tank** | 0.0015 |
| **IVC-like vs Eco-tank** | 0.0015 |
| **IVC-like vs Semi-natural** | 0.0128 |

B. Beta Diversity - Bray-Curtis (PERMANOVA, 999 permutations)

| **Source** | **Df** | **Sum of Squares** | **R^2^** | **F** | **p** |
| --- | --- | --- | --- | --- | --- |
| **Model** | 2 | 1.9078 | 0.5591 | 13.314 | 0.001 |
| **Residual** | 21 | 1.5046 | 0.4409 |  |  |
| **Total** | 23 | 3.4124 | 1 |  |  |

Pairwise PERMANOVA (FDR-adjusted)

| **Group 1** | **Group 2** | **F** | **R^2^** | **p** | **FDR** |
| --- | --- | --- | --- | --- | --- |
| Semi-natural | IVC-like | 8.4 | 0.393 | 0.001 | 0.001 |
| Semi-natural | Eco-tank | 13.15 | 0.484 | 0.001 | 0.001 |
| IVC-like | Eco-tank | 15.79 | 0.513 | 0.001 | 0.001 |

C. Homogeneity of Dispersion (PERMDISP)

| **Comparison** | **Difference** | **Lower CI** | **Upper CI** | **Adjusted p** |
| --- | --- | --- | --- | --- |
| Semi-natural vs Eco-tank | -0.113 | -0.3 | 0.072 | 0.293 |
| IVC-like vs Eco-tank | -0.082 | -0.26 | 0.095 | 0.485 |
| IVC-like vs Semi-natural | 0.03 | -0.16 | 0.219 | 0.914 |

(No significant differences in dispersion were detected.)

D. Beta Diversity - Jaccard (PERMANOVA, 999 permutations)

| **Source** | **Df** | **Sum of Squares** | **R^2^** | **F** | **p** |
| --- | --- | --- | --- | --- | --- |
| Model | 2 | 2.4622 | 0.585 | 14.799 | 0.001 |
| Residual | 21 | 1.747 | 0.415 |  |  |
| Total | 23 | 4.2092 | 1 |  |  |

**Suppl. Table S5.** Statistical analysis for Figure 6.

PERMANOVA analyses were performed using 999 permutations. Soil samples were excluded from statistical testing. Pairwise p-values were adjusted using FDR correction. Homogeneity of dispersion was assessed using betadisper and Tukey HSD post hoc testing.

A. Global PERMANOVA of gut microbiota composition (Bray-Curtis, soil excluded)

| **Source** | **Df** | **Sum of Squares** | **R^2^** | **F** | **p-value** |
| --- | --- | --- | --- | --- | --- |
| GROUP | 4 | 7.4853 | 0.5431 | 15.153 | 0.001 |
| Residual | 51 | 6.2983 | 0.4569 |  |  |
| Total | 55 | 13.7836 | 1 |  |  |

B. Pairwise PERMANOVA comparisons (Bray-Curtis, soil excluded)

| **Group 1** | **Group 2** | **F** | **R^2^** | **p-value** | **FDR-adjusted p** |
| --- | --- | --- | --- | --- | --- |
| Wild | IVC-like Mus | 7.812 | 0.303 | 0.001 | 0.001 |
| Wild | Eco-tank Mus | 3.505 | 0.132 | 0.001 | 0.001 |
| Wild | IVC-like B6 | 29.496 | 0.711 | 0.001 | 0.001 |
| Wild | Eco-tank B6 | 14.343 | 0.525 | 0.001 | 0.001 |
| IVC-like Mus | Eco-tank Mus | 8.951 | 0.224 | 0.001 | 0.001 |
| IVC-like Mus | IVC-like B6 | 27.831 | 0.582 | 0.001 | 0.001 |
| IVC-like Mus | Eco-tank B6 | 20.72 | 0.497 | 0.001 | 0.001 |
| Eco-tank Mus | IVC-like B6 | 25.37 | 0.504 | 0.001 | 0.001 |
| Eco-tank Mus | Eco-tank B6 | 15.451 | 0.373 | 0.001 | 0.001 |
| IVC-like B6 | Eco-tank B6 | 15.794 | 0.513 | 0.001 | 0.001 |

C. Homogeneity of multivariate dispersion (PERMDISP, Bray-Curtis)

| **Source** | **Df** | **Sum Sq** | **Mean Sq** | **F** | **p-value** |
| --- | --- | --- | --- | --- | --- |
| Groups | 4 | 0.2291 | 0.05727 | 4.607 | 0.004 |
| Residual | 51 | 0.634 | 0.01243 |  |  |

D. Significant Pairwise Differences (Tukey HSD)

| **Comparison** | **Adjusted p** |
| --- | --- |
| IVC-like B6 vs Eco-tank Mus | 0.00138 |

(All other pairwise comparisons *p* > 0.05)

E. Global PERMANOVA of gut microbiota composition (Jaccard, soil excluded)

| **Source** | **Df** | **Sum of Squares** | **R^2^** | **F** | **p-value** |
| --- | --- | --- | --- | --- | --- |
| GROUP | 4 | 6.6982 | 0.5549 | 15.896 | 0.001 |
| Residual | 51 | 5.3727 | 0.4451 |  |  |
| Total | 55 | 12.0709 | 1 |  |  |

F. Pairwise PERMANOVA comparisons (Jaccard, soil excluded)

| **Group 1** | **Group 2** | **F** | **R^2^** | **p-value** | **FDR-adjusted p** |
| --- | --- | --- | --- | --- | --- |
| Wild | IVC-like Mus | 6.194 | 0.256 | 0.001 | 0.001 |
| Wild | Eco-tank Mus | 5.628 | 0.197 | 0.001 | 0.001 |
| Wild | IVC-like B6 | 21.513 | 0.642 | 0.001 | 0.001 |
| Wild | Eco-tank B6 | 9.659 | 0.426 | 0.001 | 0.001 |
| IVC-like Mus | Eco-tank Mus | 9.754 | 0.239 | 0.001 | 0.001 |
| IVC-like Mus | IVC-like B6 | 29.199 | 0.593 | 0.001 | 0.001 |
| IVC-like Mus | Eco-tank B6 | 18.988 | 0.475 | 0.001 | 0.001 |
| Eco-tank Mus | IVC-like B6 | 32.353 | 0.564 | 0.001 | 0.001 |
| Eco-tank Mus | Eco-tank B6 | 15.965 | 0.38 | 0.001 | 0.001 |
| IVC-like B6 | Eco-tank B6 | 18.629 | 0.554 | 0.001 | 0.001 |

G. Homogeneity of multivariate dispersion (PERMDISP, Jaccard)

| **Source** | **Df** | **Sum Sq** | **Mean Sq** | **F** | **p-value** |
| --- | --- | --- | --- | --- | --- |
| Groups | 4 | 0.1031 | 0.02577 | 6.307 | 0.001 |
| Residual | 51 | 0.2084 | 0.00409 |  |  |

H. Significant Pairwise Differences (Tukey HSD)

| **Comparison** | **Adjusted p** |
| --- | --- |
| IVC-like B6 vs Eco-tank B6 | 0.00048 |
| IVC-like B6 vs Eco-tank Mus | 0.00088 |
| IVC-like Mus vs IVC-like B6 | 0.00201 |
| Wild vs IVC-like B6 | 0.00414 |

(All other pairwise comparisons *p* > 0.05)

I. Genotype × Housing PERMANOVA (Bray-Curtis dissimilarity; marginal tests; excluding wild state)

| **Analysis** | **R^2^** | **F** | **p-value** |
| --- | --- | --- | --- |
| Genotype × Housing | 0.091 | 7.34 | 0.001 |
| Housing (B6) | 0.513 | 15.79 | 0.001 |
| Housing (*Mus*) | 0.293 | 7.05 | 0.001 |
| Dispersion | - | 2.46 | 0.067 |

**Suppl. Table S6.** Statistical analysis for Figure 7A.

A. Pairwise PERMANOVA comparisons

Bray-Curtis dissimilarity of predicted functional profiles (999 permutations).

| **Group 1** | **Group 2** | **F** | **R^2^** | **p** | **FDR** |
| --- | --- | --- | --- | --- | --- |
| Wild | IVC | 21.804 | 0.5478 | 0.001 | 0.0015 |
| Wild | Eco-tank | 2.558 | 0.1001 | 0.051 | 0.051 |
| IVC | Eco-tank | 11.356 | 0.2681 | 0.001 | 0.0015 |

B. Overall PERMANOVA model

| **Source** | **Df** | **Sum of Squares** | **R^2^** | **F** | **p** |
| --- | --- | --- | --- | --- | --- |
| Model | 2 | 0.39596 | 0.3513 | 9.748 | 0.001 |
| Residual | 36 | 0.73115 | 0.6487 |  |  |
| Total | 38 | 1.12711 | 1 |  |  |

**Suppl. Table S7.** Statistical analysis for Figure 8A.

A. Pairwise PERMANOVA comparisons

Bray-Curtis dissimilarity of predicted functional profiles (999 permutations).

| **Group 1** | **Group 2** | **F** | **R^2^** | **p** | **FDR** |
| --- | --- | --- | --- | --- | --- |
| Semi-natural | IVC-like | 8.4 | 0.393 | 0.003 | 0.003 |
| Semi-natural | Eco-tank | 13.15 | 0.484 | 0.001 | 0.002 |
| IVC-like | Eco-tank | 15.79 | 0.513 | 0.001 | 0.002 |

B. Overall PERMANOVA model

| **Source** | **Df** | **Sum of Squares** | **R^2^** | **F** | **p** |
| --- | --- | --- | --- | --- | --- |
| Model | 2 | 1.908 | 0.559 | 13.314 | 0.001 |
| Residual | 21 | 1.505 | 0.441 |  |  |
| Total | 23 | 3.412 | 1 |  |  |

**Supplemental Figure S1.** Experimental design for microbiota conditioning strategies in Wild *Mus* *musculus* and C57BL/6 mice, and vaccination workflow in eco-tank. (A) Schematic overview of the experimental timeline and housing conditions for wild-derived mice. Following capture (Wild) and initial laboratory acclimation under standard chow diet for 14 days (IVC-like cage), mice were assigned to different housing and dietary enrichment conditions, including IVC-like cages, soil bedding, wild mouse diet, or combined semi-natural cages (Day 14-24). On Day 24, enriched groups were transferred to a large semi-natural enclosure (Eco-tank), while chow-only controls remained in standard laboratory cages. Microbiome analyses were conducted longitudinally through Day 52. Dashed lines indicate sampling time points. (B) C57BL/6 mice were assigned to one of three housing and microbiota exposure conditions (IVC-like; semi-natural cage; eco-tank). Arrows indicate longitudinal experimental follow-up under each condition (28 days). (C) C57BL/6 mice were housed in eco-tank. For vaccination or infection studies, mice were temporarily removed from the Eco-tank and housed in semi-natural cages for 24-48 hours to allow administration of vaccine or infection under controlled laboratory conditions. Following vaccination, mice were returned to their original Eco-tank to maintain microbiota conditioning. For infection experiments, mice were euthanized at 24 hours post-infection for quantification of bacterial burden (colony-forming units, CFU) in lungs. Figure was generated by using BioRender.

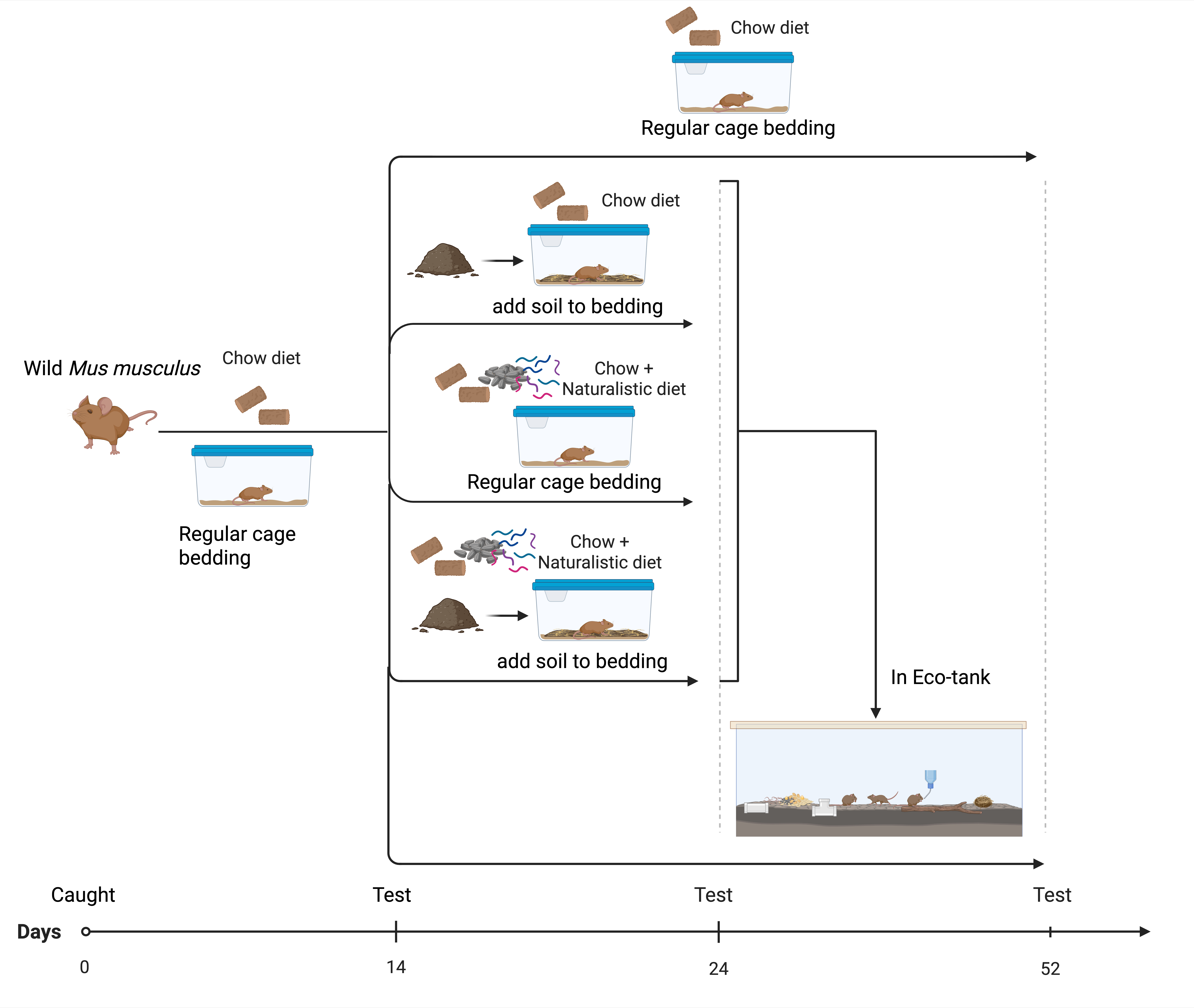

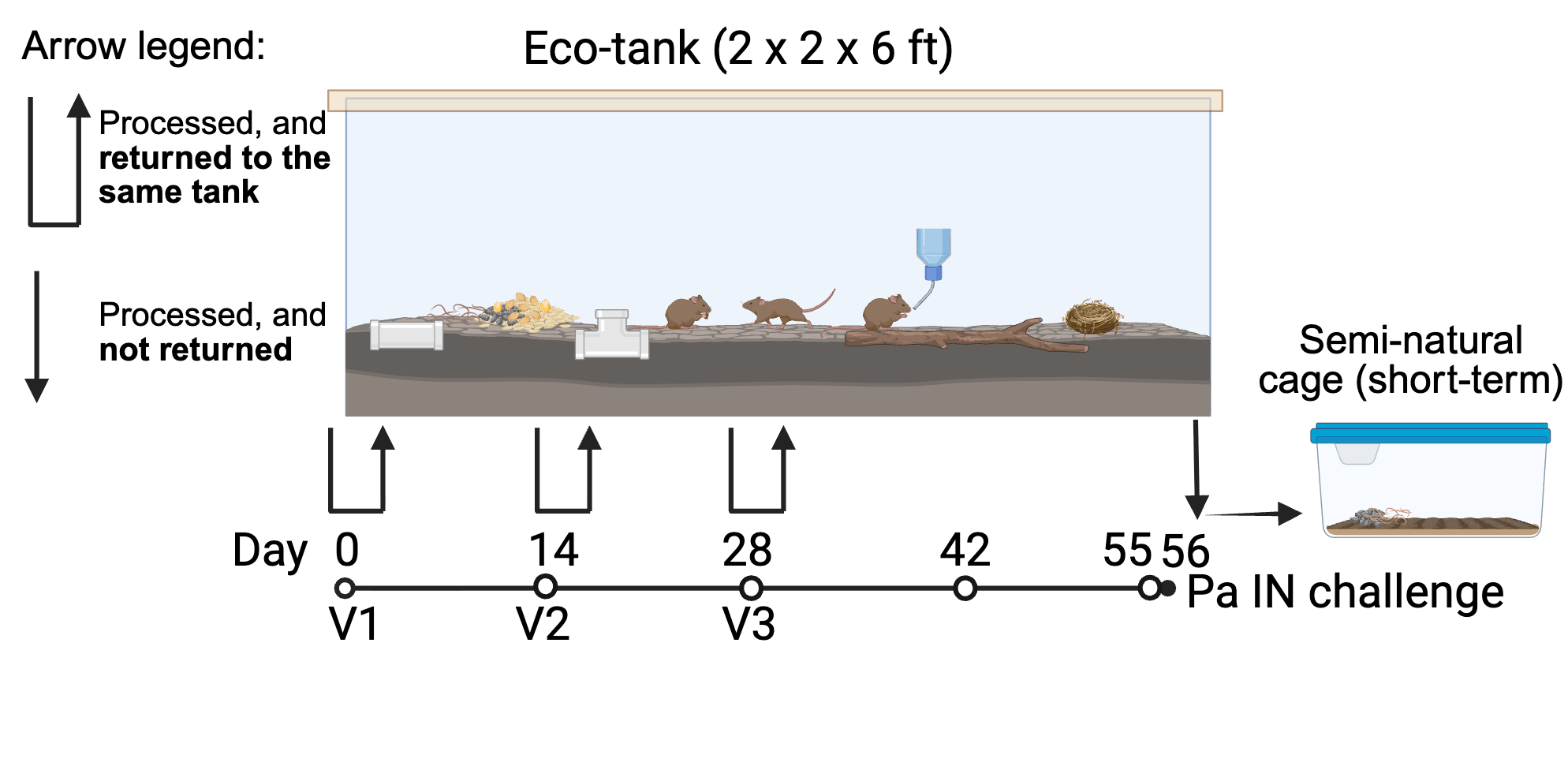

**B**

**C**

**A**

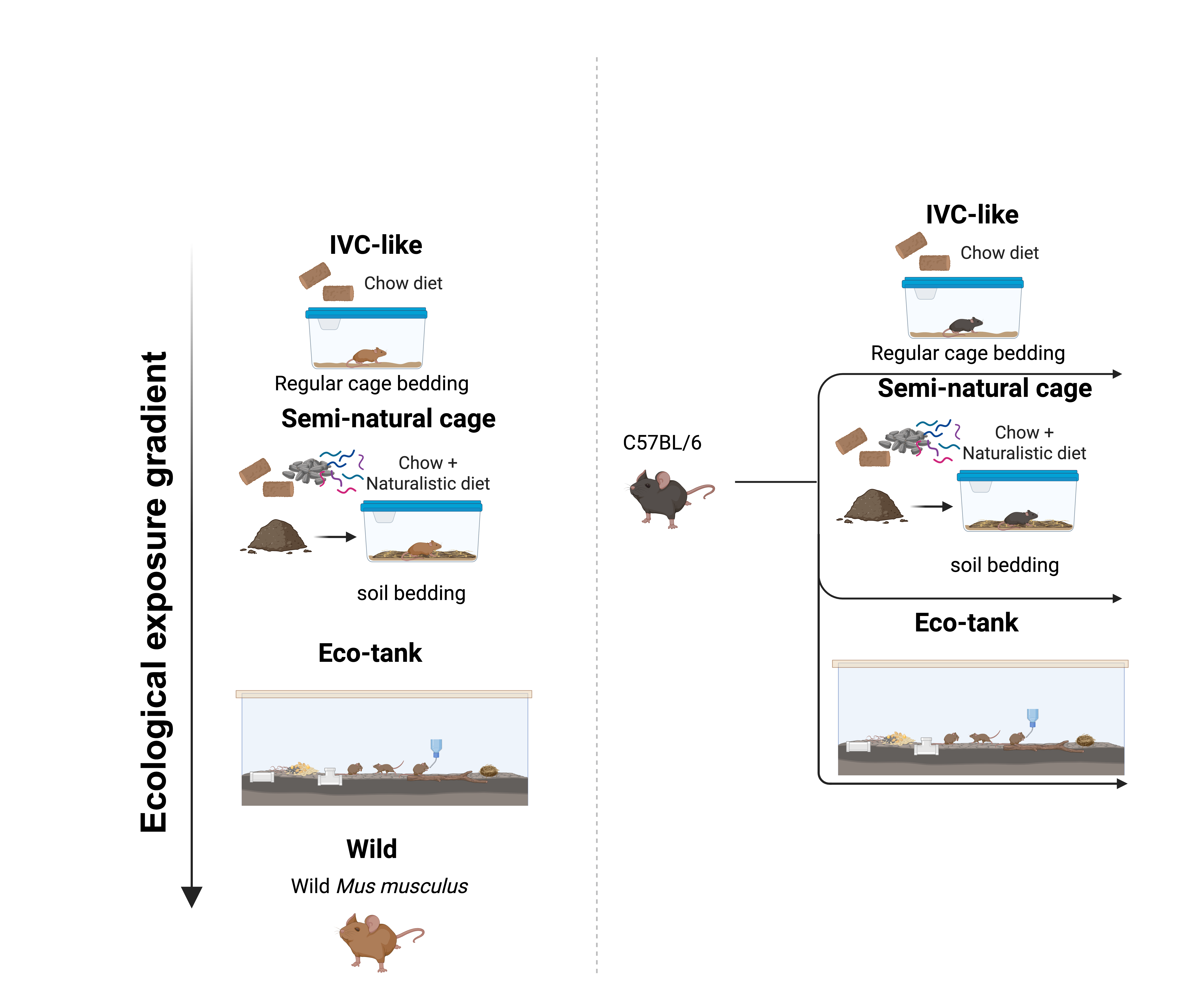

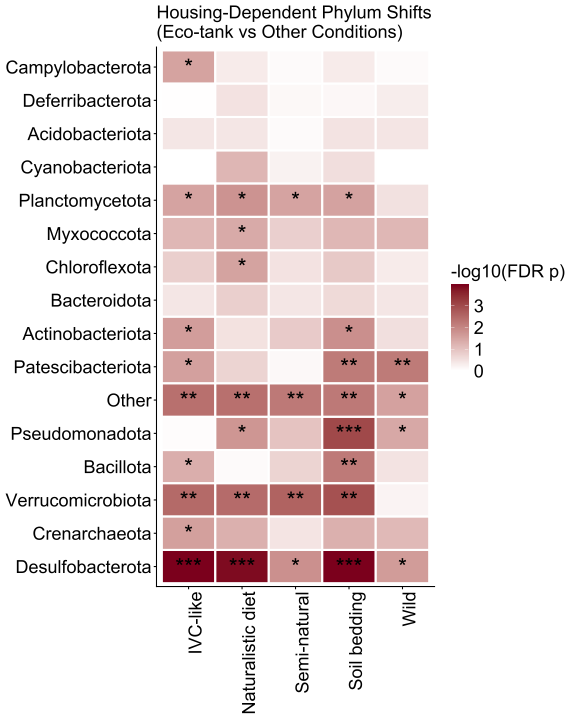

**C**

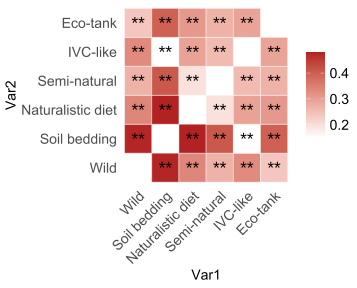

**A**

**B**

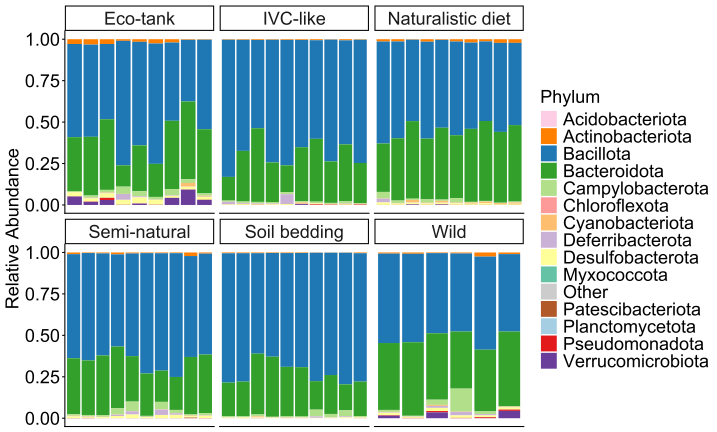

**Supplemental Figure S2.** (A) Pairwise effect size heatmap of housing-dependent microbiota restructuring in wild *Mus*. Heatmap summarizing pairwise PERMANOVA effect sizes (R^2^) for ASV-level Bray-Curtis dissimilarity across housing conditions. Each tile represents the proportion of variance explained (R^2^) for the comparison between two groups. Warmer colors indicate stronger compositional separation. Statistical significance was determined using permutation-based PERMANOVA with false discovery rate (FDR) correction for multiple comparisons. Asterisks indicate adjusted p-values (**FDR < 0.01). Diagonal entries represent within-group comparisons and are shown as zero. (B) Phylum-level taxonomic composition of gut microbiota across experimental conditions at the indicated time points. Bars represent individual relative abundance of each sample. (C) Heatmap depicting housing-associated differences in phylum-level relative abundance compared to eco-tank conditions. Color intensity represents −log10(FDR-adjusted *p*-value) from pairwise comparisons. Asterisks denote significance thresholds (* FDR < 0.05, ** FDR < 0.01, *** FDR < 0.001).

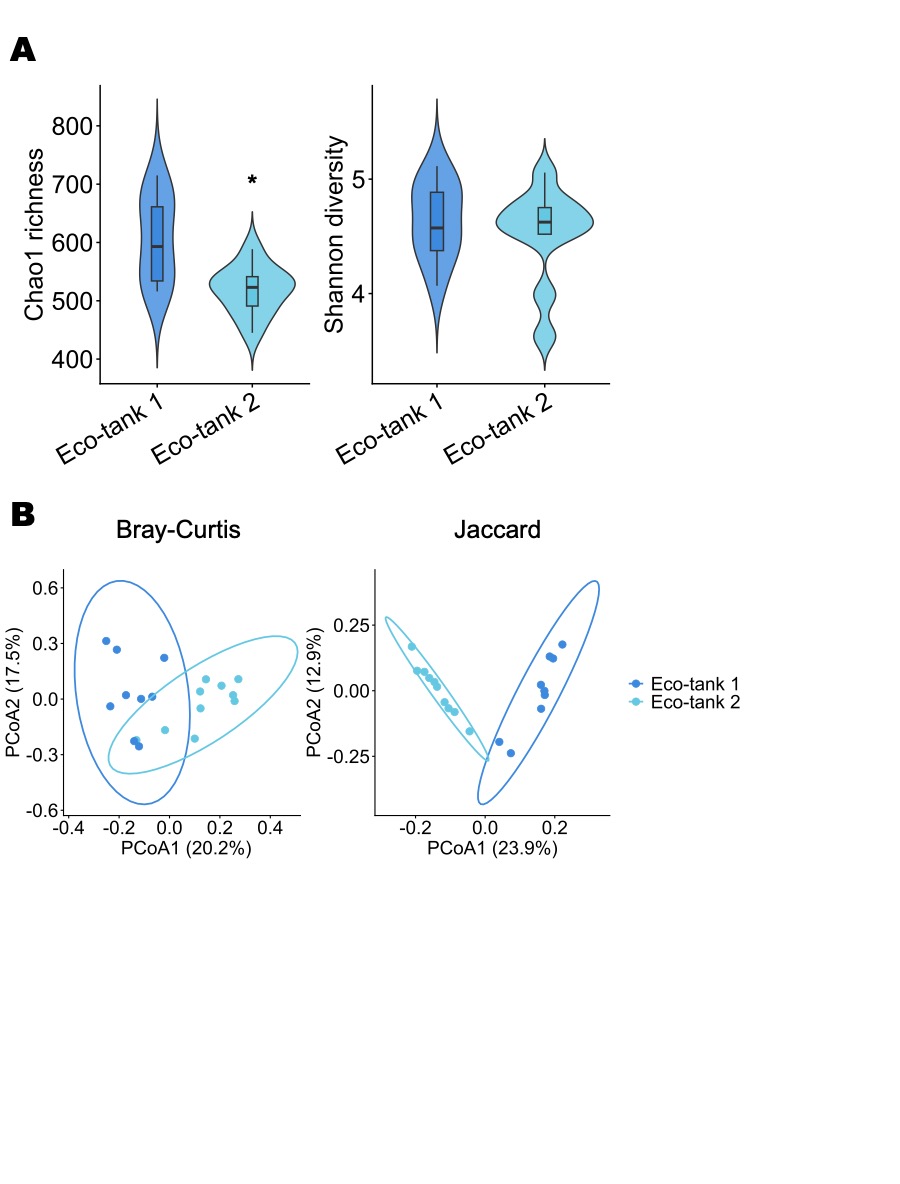

**Supplemental Figure S3.** Independent Eco-tanks yield broadly similar gut microbiota configurations. (A) Alpha diversity of gut microbiota in wild-derived *Mus musculus* housed in two independently established Eco-tanks (Eco-tank 1: blue and Eco-tank 2: cyan). Chao1 richness (left; **p* = 0.02474) and Shannon diversity (right; *p* = 0.6534) are shown as violin plots with embedded boxplots. (B) Principal coordinates analysis (PCoA) of beta diversity based on Bray-Curtis (left; PERMANOVA: R^2^ = 0.16, *p* = 0.001; PERMDISP: *p* = 0.83) and Jaccard (right; PERMANOVA: R^2^ = 0.22, *p* = 0.001; PERMDISP: *p* = 0.155) dissimilarities. Each point represents an individual mouse; ellipses denote 95% confidence intervals. Substantial overlap was observed between Eco-tank 1 and Eco-tank 2 communities.

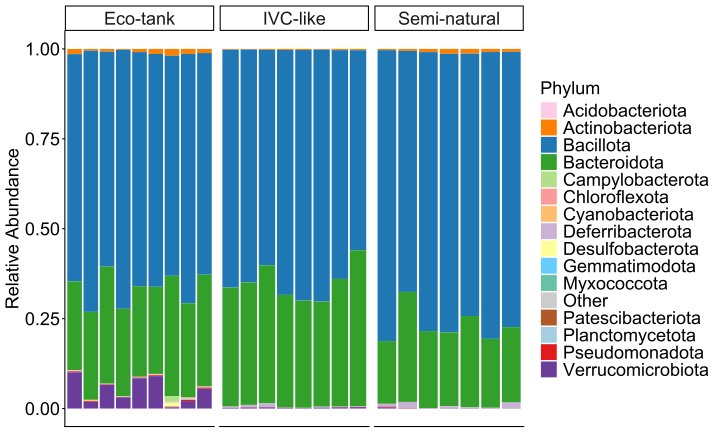

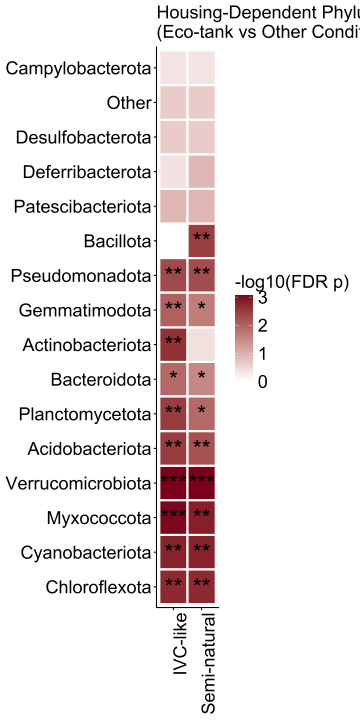

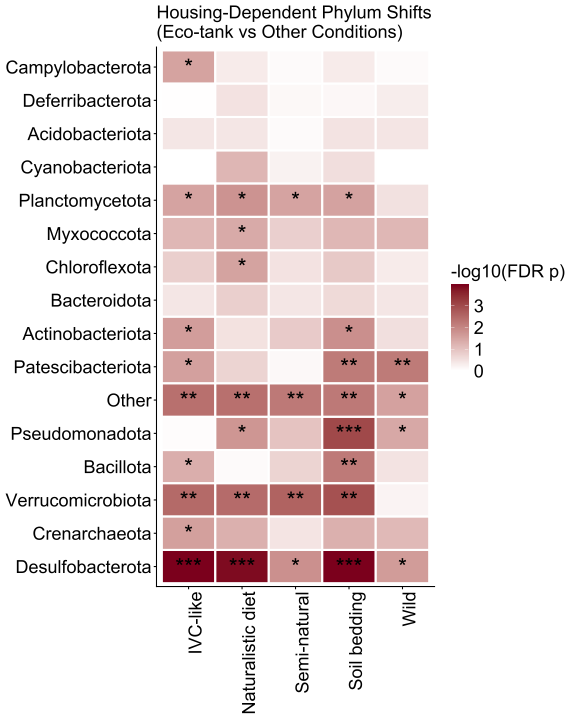

**A**

**B**

**Supplemental Figure S4.** (A) Phylum-level taxonomic composition of gut microbiota across experimental conditions at the indicated time points. Bars represent individual relative abundance of each sample. (B) Heatmap depicting housing-associated differences in phylum-level relative abundance compared to eco-tank conditions. Color intensity represents −log10(FDR-adjusted p-value) from pairwise comparisons. Asterisks denote significance thresholds (* FDR < 0.05, ** FDR < 0.01, *** FDR < 0.001).

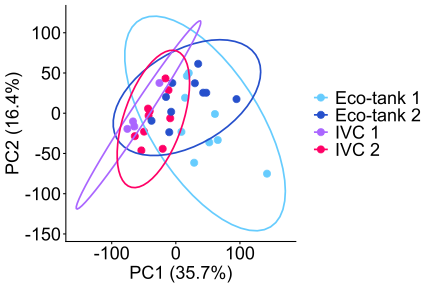

**Supplemental Figure S5.** Principal component analysis (PCA) of predicted functional profiles generated using TAX4FUN2 from ASV-level data. Functional abundances (KEGG Ortholog level) were normalized to relative abundance and scaled prior to analysis. Each point represents one fecal sample. Percent variance explained by each principal component is indicated on the axes. Group differences in overall functional composition were assessed using Bray-Curtis dissimilarity. Ellipses indicate 95% confidence intervals.

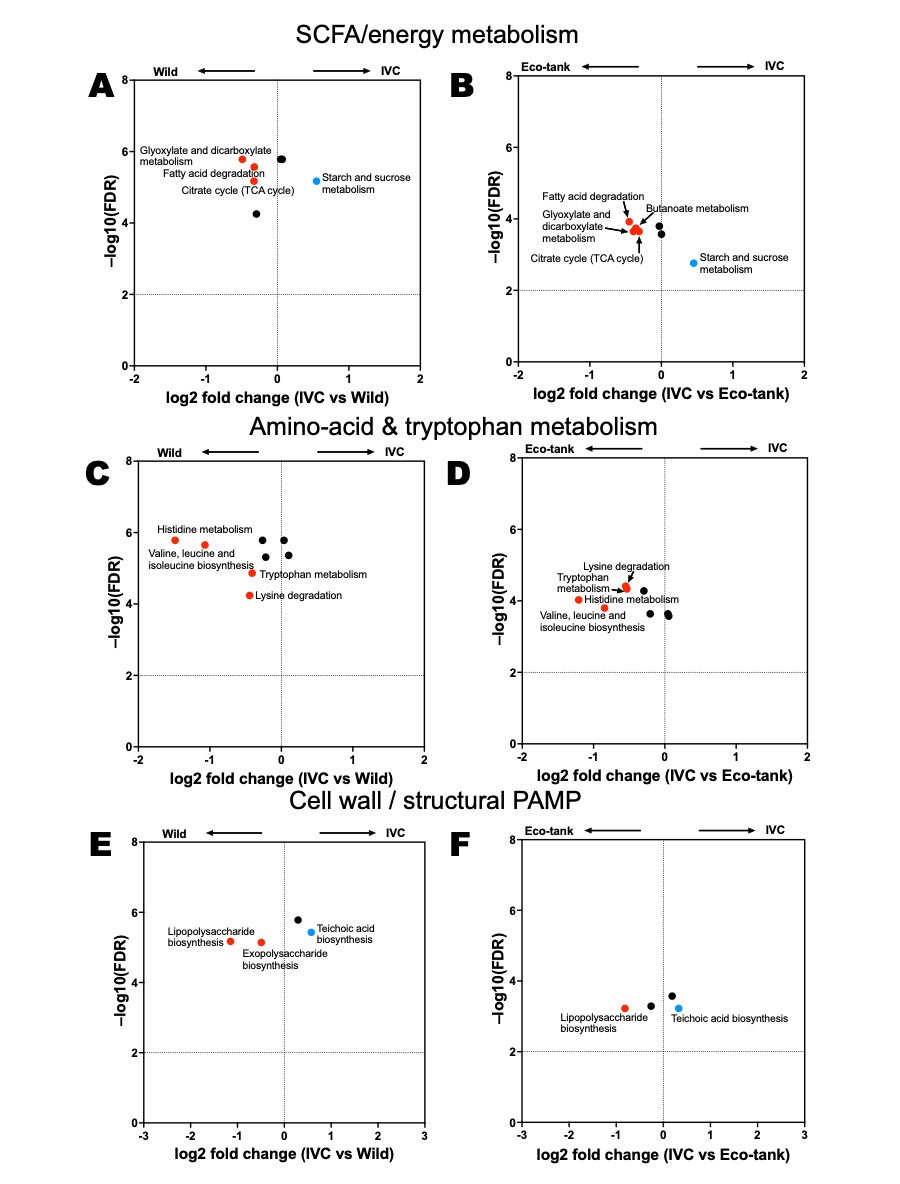

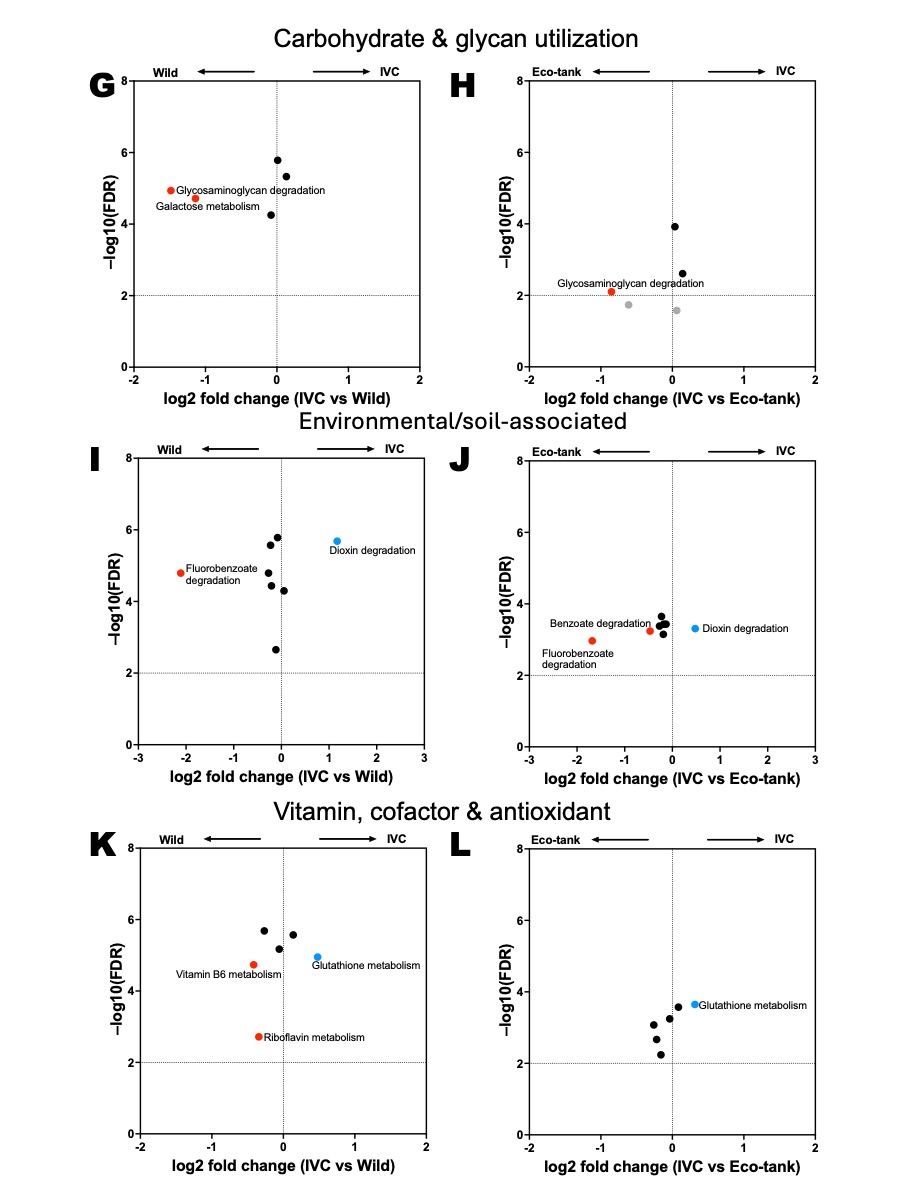

**Supplemental Figure S6.** Eco-tank housing selectively preserves wild-associated microbial functional pathways. Volcano plots showing differential abundance of predicted KEGG pathways (Tax4Fun2) comparing IVC-housed mice to Wild mice (left panels) or Eco-tank housed mice (right panels). Functional categories include SCFA and energy metabolism (A-B), amino acid and tryptophan metabolism (C-D), cell wall and structural PAMP biosynthesis (E-F), carbohydrate and glycan utilization (G-H), environmental/soil-associated metabolism (I-J), and vitamin, cofactor, and antioxidant pathways (K-L). Each point represents a KEGG pathway. The x-axis indicates log2 fold change (IVC vs Wild or IVC vs Eco-tank), and the y-axis indicates -log10(FDR). Colored pathways met an FDR threshold < 0.01. Red points indicate pathways enriched in wild or Eco-tank mice relative to IVC (log2FC <−0.3), and blue points indicate pathways enriched in IVC-like mice (log2FC > 0.3).
